# Pervasive effects of *Wolbachia* on host body mass and metabolic rate

**DOI:** 10.64898/2026.09.09.750401

**Authors:** Jason M. Graham, H. Arthur Woods, Brandon S. Cooper, Michael T.J. Hague

**Affiliations:** Department of Mathematics, University of Scranton, 800 Linden Street Scranton, PA 18510; Division of Biological Sciences, University of Montana, 32 Campus Dr. Missoula, MT 59812; Biology Department University of Scranton 800 Linden Street Scranton, PA 18510

**Keywords:** host-microbe interaction, endosymbiosis, *Drosophila*, *w*Mel, metabolic rate

## Abstract

Metabolic rate is a widely variable trait that reflects organismal energy expenditure. At least half of all arthropod species carry maternally transmitted *Wolbachia* bacteria, but it is poorly understood how these endosymbionts affect host physiology and fitness. We tested how two central host traits, body mass and standard metabolic rate (SMR), are altered by *Wolbachia* for nine diverse *Drosophila* fly species and 13 co-occurring *Wolbachia* variants diverged roughly 150 million years. Interspecific trait variation was structured according to the host phylogeny, but within each host system, *Wolbachia* almost always altered body mass and/or SMR. *Wolbachia* often caused increases or decreases in body mass that, in some cases, also led to mass-dependent effects on whole-organism SMR. Other *Wolbachia* altered mass-adjusted SMR. These variable effects are plausibly explained by complex interactions among *Wolbachia* and host genomes, as documented for other traits (e.g., *Wolbachia*-induced cytoplasmic incompatibility). Endosymbiont effects on body mass and metabolic rate likely have cascading consequences for host biology and evolution, because these central traits are correlated with many aspects of host physiology, behavior, and life history. This also applies to hosts transinfected with *Wolbachia* from *Drosophila* to reduce the impacts of pathogens and pests on multiple continents.

## INTRODUCTION

Metabolic rate is a measure of the energy that organisms expend to sustain life, integrating many aspects of organismal physiology, behavior, and life history. Metabolic rates vary widely among individuals and species (Burton et al. 2011; Konarzewski and Książek 2013; White and Kearney 2013; Auer et al. 2017), with implications for components of fitness, including growth, survival, behavior, and reproduction (Glazier 2005; Biro and Stamps 2010; Burton et al. 2011; Pettersen et al. 2016; Pettersen et al. 2018; Glazier 2022; White and Marshall 2023). Resting metabolic rate and other baseline measures of metabolism characterize the minimal energy expenditure required for self-maintenance (Stearns 1998; Burton et al. 2011; Pontzer and McGrosky 2022; White et al. 2022). In arthropods and other ectotherms, standard metabolic rate (SMR) represents the lowest rate of metabolism in an inactive, post-absorptive, nonreproductive adult, measured at a particular temperature. SMR is highly variable in insects and other arthropods (e.g., Addo-Bediako et al. 2002; Nespolo et al. 2003; Messamah et al. 2017; Careau et al. 2019; Somjee et al. 2021; Leahy et al. 2025) and identifying the underlying sources of this variation is central to understanding broadscale patterns of energy expenditure.

A key source of variation in metabolic rate is body size, which generally scales hypometrically with metabolic rate, both within and among species (Addo-Bediako et al. 2002; Glazier 2005; Niven and Scharlemann 2005; Chown et al. 2007; White et al. 2007; Kolokotrones et al. 2010; White and Kearney 2011; White and Kearney 2013; White et al. 2019; Glazier 2022; Harrison et al. 2022); although, the relationship between body mass and metabolic rate is generally less strong for ectotherms (White et al. 2007). After accounting for body mass, both intrinsic factors (e.g., sex: Tomlinson and Phillips 2015; Arnqvist et al. 2017; Videlier et al. 2019; mitochondrial function: Speakman et al. 2004; Tieleman et al. 2009; Hoekstra et al. 2013; Hoekstra et al. 2018; Matoo et al. 2019) and extrinsic factors (e.g., temperature: Gillooly et al. 2001; Addo-Bediako et al. 2002; Nespolo et al. 2003; Lachenicht et al. 2010; Alton et al. 2017) contribute to mass-adjusted metabolic rates. Even after accounting for these factors and controlling for phylogeny (Addo-Bediako et al. 2002; Chown et al. 2007; Leahy et al. 2025), there remains much unexplained variation in metabolic rates (Burton et al. 2011; Videlier et al. 2019; Glazier 2022; Harrison et al. 2022; Leahy et al. 2025).

For insects and other arthropods, interactions with heritable microbes may contribute to metabolic-rate variation. Insects host a diversity of endosymbionts that alter central aspects of host biology and fitness (Moran et al. 2008; Werren et al. 2008; Feldhaar 2011; McFall-Ngai et al. 2013; McCutcheon et al. 2019; Hoffmann and Cooper 2024). Endosymbionts may require additional energy expenditures by hosts (Evans et al. 2009; Clavé et al. 2022), considering they interact with aspects of host nutrition (Koga et al. 2003; Brownlie et al. 2009; Kwak et al. 2025; Lindsey et al. 2025; Preuss et al. 2026), immune function (Oliver et al. 2003; Hedges et al. 2008; Teixeira et al. 2008; Jaenike et al. 2010; Nichols et al. 2021), and thermoregulation (Russell and Moran 2005; Brumin et al. 2011; Hague, Caldwell, et al. 2020). Endosymbionts are also maternally co-inherited with mitochondria in the cytoplasm (Ballad et al. 1996; Bright and Bulgheresi 2010; Richardson et al. 2012; Russell et al. 2019; Camus et al. 2022). Mitochondrial function and cytonuclear interactions with the nuclear genetic background impact SMR and other metabolic traits (e.g., Arnqvist et al. 2010; Montooth et al. 2010; Mossman et al. 2016); however, it is generally unknown how metabolic rate is affected when endosymbionts are present in the cytoplasm (but see Evans et al. 2009).

Here, we test the hypothesis that endosymbionts alter host metabolic rate. We examined variation in body mass and metabolic rates (SMR) in diverse *Drosophila* fly species and then tested how *Wolbachia* endosymbionts affect these central organismal traits. Maternally transmitted *Wolbachia* infect at least half of arthropod species (Weinert et al. 2015) and many insects (Werren et al. 2008; Zug and Hammerstein 2012). *Wolbachia* are usually facultative and not required for host survival (Hoffmann and Cooper 2024; Hoffmann and Cooper 2025), with population frequencies varying over space and time (Hoffmann et al. 1998; Kriesner et al. 2013; Hamm et al. 2014; Cooper et al. 2017; Hague, Mavengere, et al. 2020; Ravikanthachari et al. 2026). Body mass and metabolic rates are widely studied in arthropods, including in *Drosophila* (e.g., Addo-Bediako et al. 2002; Chown et al. 2007; Jumbo-Lucioni et al. 2010; Messamah et al. 2017; Videlier et al. 2019; Videlier et al. 2021), but the effects of *Wolbachia* are generally not considered.

*Wolbachia* cells are found throughout host tissues, including gonads, fat body, gut, salivary glands, hemocytes, and Malpighian tubules (reviewed in Pietri et al., 2016). *Wolbachia* localization to reproductive tissues mediates maternal transmission (Russell et al. 2019; Porter and Sullivan 2023; Radousky et al. 2023; Hague et al. 2024) and effects on host reproduction (cytoplasmic incompatibility; Shropshire et al. 2020; Shropshire et al. 2022); however, the physiological consequences of localization in somatic tissues are largely unknown (Hosokawa et al. 2010; Moriyama et al. 2015; Hague, Caldwell, et al. 2020; Porter and Sullivan 2023). Understanding how *Wolbachia* impact central organismal traits like metabolic rate is particularly important, because little is known about the fitness effects that underlie *Wolbachia* spread in nature (Hoffmann et al. 1990; Kriesner and Hoffmann 2018; Ross et al. 2019; Hoffmann and Cooper 2024; Graham et al. 2025); although effects on fecundity, virus blocking, and other traits remain candidates (Weeks et al. 2007; Brownlie et al. 2009; Cogni et al. 2021). It is also critical for improving *Wolbachia*-based biocontrol programs to prevent human diseases (Hoffmann et al. 2011; Walker et al. 2011; Ross et al. 2019; Utarini et al. 2021; Lenharo 2023; Velez et al. 2023) and agricultural pests (Nikolouli et al. 2018; Gong et al. 2020), where transinfected hosts experience fitness costs (reviewed in Ross et al. 2019).

We quantified body mass and SMR for 3,102 adult flies from nine different *Drosophila* species, with and without their naturally occurring *Wolbachia* strains, to test whether endosymbionts alter host metabolic rates. We examined the effects of 13 divergent *Wolbachia* variants (Figure 1), many of which have recently switched hosts via horizontal and introgressive transfer (Turelli et al. 2018; Cooper et al. 2019; Shropshire et al. 2026). We documented pervasive—but variable—effects of *Wolbachia* on body mass and metabolic rate in almost all *Wolbachia*-host systems assayed. In some cases, *Wolbachia* effects on body mass altered whole-organism SMR due to the scaling relationship between body mass and metabolic rate. Other *Wolbachia* variants altered mass-adjusted SMR after controlling for body mass. The widespread, variable effects we observed may be explained by novel cytonuclear interactions that have resulted from rapid *Wolbachia* host switching. Our findings imply that facultative endosymbionts are an important, unexplored source of variation in metabolic rates and related organismal traits.

**Figure 1.**
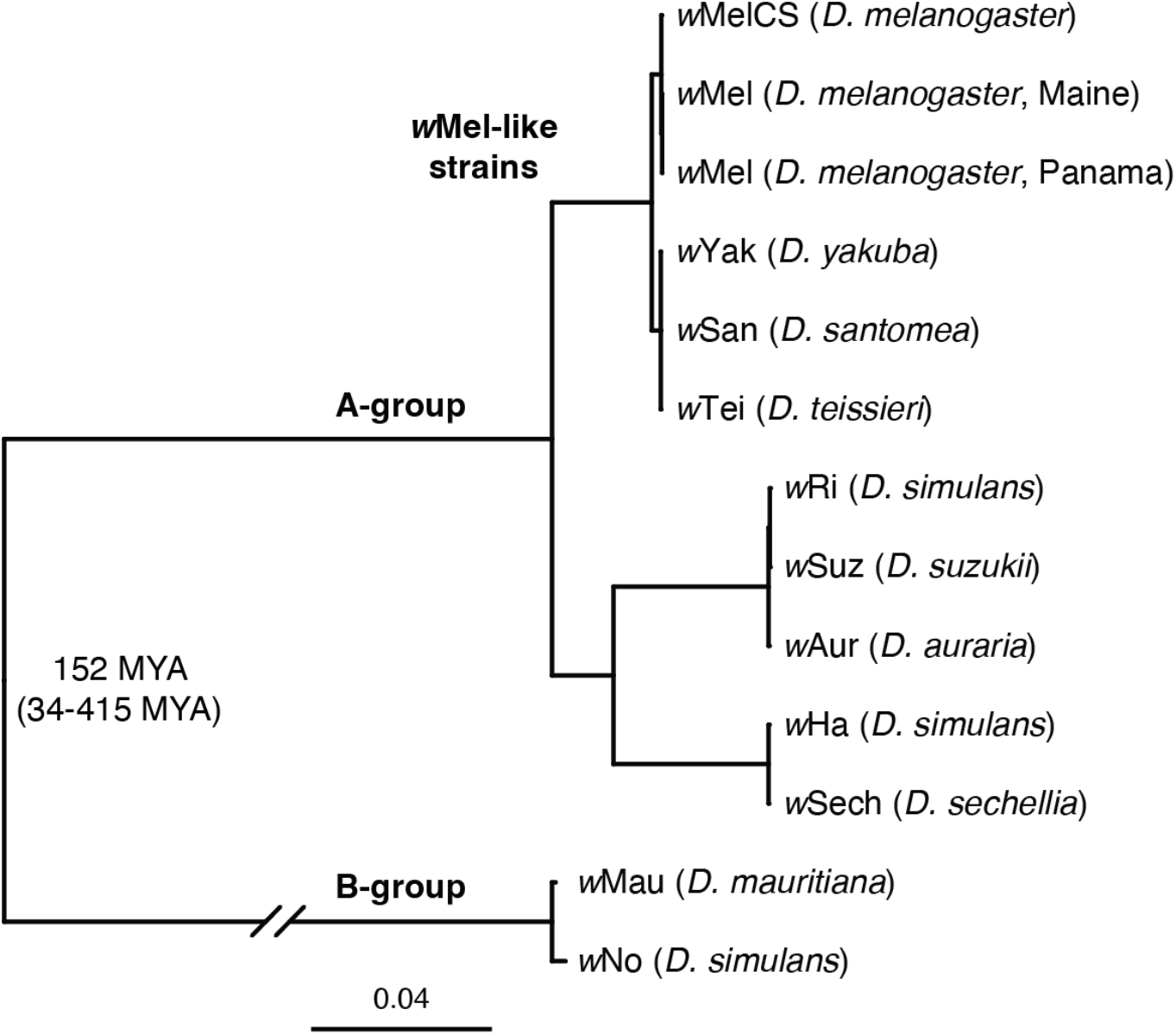
Divergent *Wolbachia* strains found in *Drosphila* host species. Estimated Bayesian phylogram of A-and B-group *Wolbachia* strains using 211 single copy genes of identical length spanning a total of 178,569 bp. The estimated divergence date with confidence intervals for A-and B-groups is reproduced from Shropshire et al. (2026). Naturally occurring *Drosophila* host species are listed in parentheses.

## MATERIALS AND METHODS

### Flies

We quantified body mass and SMR for *Wolbachia*-negative and -positive flies from nine different *Drosophila* species spanning roughly 22 million years of host evolution (Figure 2, Supplemental Table S1) (Suvorov et al. 2022; Shropshire et al. 2026). For two of the fly species, *D. melanogaster* and *D. simulans*, we tested multiple genotypes carrying different *Wolbachia* variants, which included *w*Mel-*D. melanogaster* genotypes from northern and southern latitudes (Maine, USA and Panama City, Panama, respectively) and *D. simulans* lines carrying *w*Ri, *w*Ha, and a co-infection of *w*Ha and the B-group *w*No strain (Figure 1) (O’Neill and Karr 1990; Mercot et al. 1995; Rousset and Solignac 1995; James et al. 2002). In total, the dataset included 13 different *Wolbachia*-*Drosophila* host systems. First, we analyzed *Wolbachia*-negative flies from each genotype (*N* = 1,544) to quantify interspecific patterns of variation in body mass and SMR in a comparative framework, absent of any putative effects from *Wolbachia*. Next, we incorporated *Wolbachia*-positive individuals (*N* = 1,558) into focused analyses of each genotype to evaluate how *Wolbachia* alter body mass and SMR within each *Wolbachia*-host system (Supplemental Figures S1, S2).

**Figure 2.**
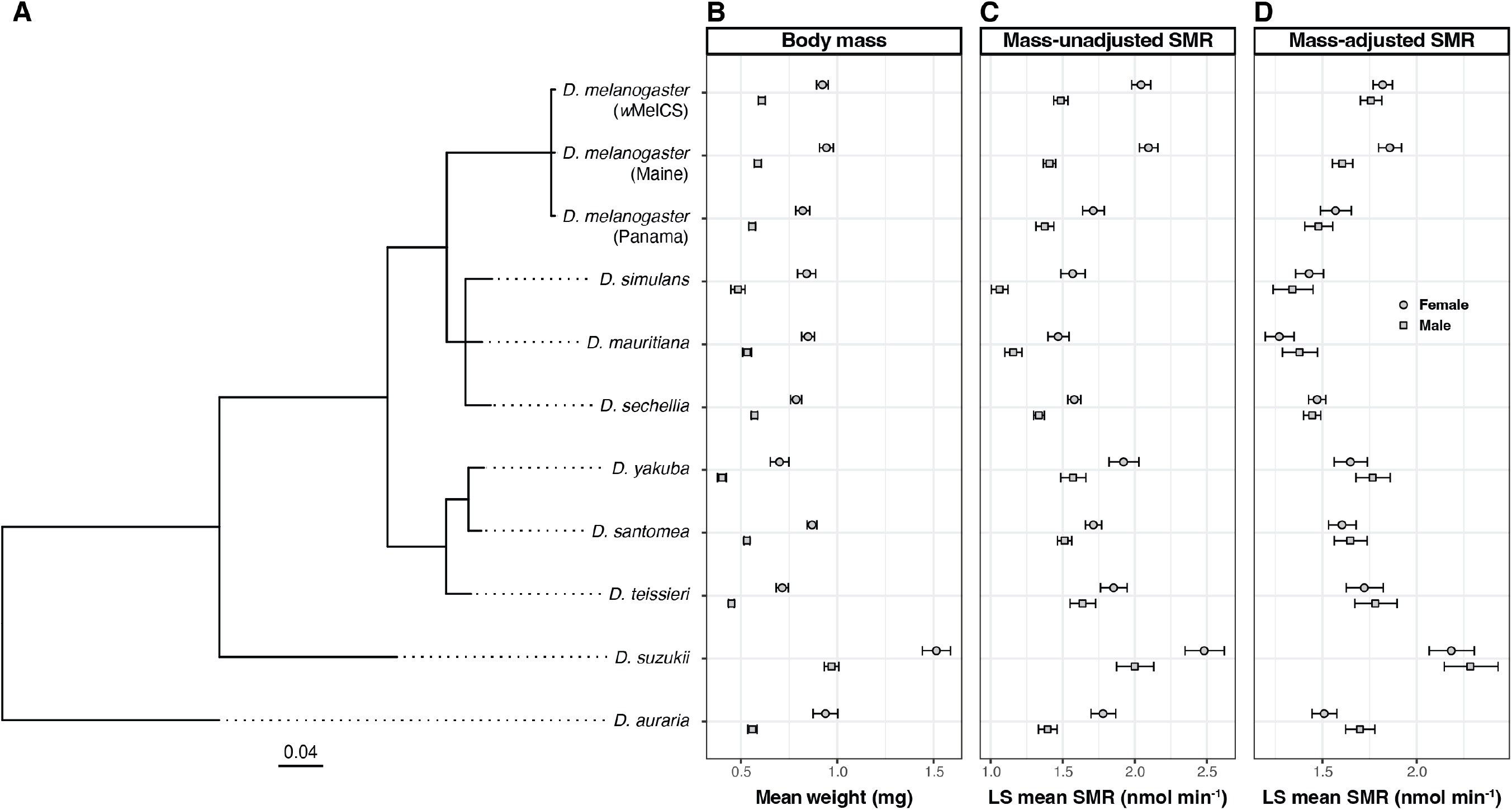
Body mass and SMR estimates from *Wolbachia*-negative flies show strong evidence of phylogenetic signal. **(A)** Estimated Bayesian phylogram of *Drosophila* host species using 20 conserved single-copy genes. **(B)** Mean body mass (± 95% CIs) of *Wolbachia*-negative flies for each sex of each host genotype. **(C)** Least square (LS) mean mass-unadjusted SMR (± 95% CIs) of *Wolbachia*-negative flies for each sex of each host genotype. LS mean values are extracted from genotype-specific LMMs that account for fly activity, experiment start time, flow rate in the chamber, water vapor, temperature, and light intensity. **(D)** LS mean mass-adjusted SMR (± 95% CIs) of *Wolbachia*-negative flies for each sex of each host genotype. LS mean values are extracted from genotype-specific LMMs that account for body mass, in addition to the nuisance variables listed in panel C. *Wolbachia*-negative data from the *w*Ri-*D. simulans* genotype are shown for the *D. simulans* tip on the phylogeny.

In an earlier study, we analyzed effects of *Wolbachia* on locomotor activity of the flies in this study (Hague et al. 2021). As described in Hague et al. (2021), *Wolbachia*-positive flies were treated with tetracycline to generate paired *Wolbachia*-negative versions of each host genotype (Cooper and Shropshire 2024). The absence of *Wolbachia* was confirmed with qPCR. The gut microbiome of the tetracycline-cleared flies was then reconstituted by rearing them on food in which *Wolbachia*-positive males of the same genotype had fed and defecated for the prior 48 hours. Extra care was taken to avoid any potential detrimental effects of the antibiotic treatment by giving the flies a minimum of three generations after the tetracycline treatment before experiments (Ballard and Melvin 2007). Experiments took place many generations after the tetracycline treatment.

All flies were reared at 25°C under a 12L:12D light cycle (Percival Model I-36LL) on a standard food diet (Hague, Caldwell, et al. 2020; Hartman et al. 2024; Wheeler et al. 2024). Each day, we collected a batch of female and male virgins from the paired *Wolbachia*-negative and - positive lines of a single genotype. The four treatment groups (*Wolbachia*-negative females, - negative males, -positive females, -positive males) were maintained separately in isolation at 25°C for 3 to 5 days prior to flowthrough respirometry experiments.

### Flowthrough respirometry

We measured metabolic rates of individual flies using a differential CO_2_ analyzer (LI7000; Li-Cor Biosciences) and 16-chamber flowthrough respirometry and data acquisition system (MAVEn, Sable Systems International). The MAVEn incorporates a flow-distribution manifold, a main board (flow measurement, regulation, and control plus data acquisition and signal processing), and an activity board (sensors for activity, ambient temperature, humidity, and light intensity). The experiments were conducted at room temperature, and the readings from the activity board were used to account for any minor fluctuations in temperature, humidity, etc. in downstream analyses.

The CO_2_ analyzer was calibrated weekly with CO_2_-free air and 20 ppm CO_2_ in N_2_ (Norco). The baseline flow rate of gas was set to 35 ml min^-1^. Approximately equivalent flow rates in the experimental channels (chambers 1-16) are maintained by means of matched flow resistances based on micro-orifice flow restrictors. During normal operation, a constant stream of dry CO_2_-free is directed into the reference cell (A) of the CO_2_ analyzer and then the airstream is humidified by directing it through Nafion tubing (Chemours) submerged in distilled water. The re-humidified airstream is then fed into the flow-distribution manifold, where it is split into 17 streams (one for each of the 16 animal chambers and one for the baseline). Each of the 16 animal chambers is a 2.4 ml volume tube. A mass flow meter on the MAVEn’s main board measures the actual flow rate of each airstream. Finally, the airflow is directed into the measurement cell (B) of the CO_2_ analyzer, which records the fractional CO_2_ concentration in excurrent air (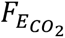).

Each day, we selected a batch of flies from a single genotype and randomly assigned the individual 3-to 5-day-old flies from each treatment group to one of the 16 animal chambers. The flies were gently aspirated into each chamber without anesthesia. Chambers were placed on the MAVEn activity board that uses infrared light (invisible to flies) to monitor animal activity in each chamber, sampled at 1 Hz. Metabolic measurements were performed between the hours of 09:00 and 16:00 during the day. We set the MAVEn to measure each chamber for 5 min (dwell time) with baseline measurements (5 min) taken every 2 chambers (i.e., interleave ratio = 2), resulting in a 120-min cycle to measure all 16 chambers. In each run, we allowed the system to complete two complete cycles, so that metabolic data for each fly was recorded twice (i.e., a minimum of 240 min). We monitored real-time flow rates to check for leaks, and chambers with erratic or unexpectedly low flows were removed prior to analysis. Immediately following each experiment, flies were anesthetized with CO_2_ and their (wet) weights were measured using an analytical balance (Mettler Toledo). Chambers were washed with water and detergent after each run.

### Data processing

Data were extracted and analyzed using R (R Core Team 2023). The CO_2_ trace was corrected for drift using the interleaving baseline means. We modeled lines between the means of each interleaving baseline measurement and then, for each sampling timepoint, subtracted the baseline estimate from the actual measurement. We also corrected for the 3-sec lag between the activity channels and the CO_2_ trace. The raw activity channels (one per chamber) were transformed into absolute difference sums (ADS). ADS was calculated by first calculating the cumulative sum of the absolute difference between consecutive activity readings and then calculating the slope of cumulative activity versus time (Videlier et al. 2019; Hague et al. 2021; Videlier et al. 2021). We discarded the initial 30 min of data from each run to avoid artifacts from the initial period during which flies adjust to the chambers.

Raw measures of CO_2_ (ppm) were converted to molar rates of CO_2_ production 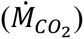 using the known flow rate (l min^-1^) and the Ideal Gas Law:

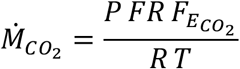

where 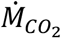 is the rate of CO_2_ production of individual flies (mol min^-1^), *P* is the pressure (1 atm), *FR* is the flow rate (l min^-1^), 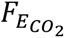 is the fractional CO_2_ concentration in the excurrent air, *R* is the gas constant (0.08206 l atm K^-1^ mol^-1^), and *T* is temperature (K). For ease of viewing, 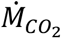 values were converted to nmol min^-1^ by multiplying by 10^9^.

The first 2 min of each metabolic measurement was discarded to allow the system to fully equilibrate after switching to a new chamber. From the remaining 3 min of each measurement period, we extracted the lowest 20-sec continuous bout of 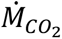 and calculated the average to generate an estimate of SMR. In addition to the average of the lowest 20-sec, we also extracted the start time of the SMR recording, the average flow rate (ml min^-1^), water vapor pressure (ppt), temperature (°C), light intensity (lux), and activity (ADS) during the recording. While we previously found that *Wolbachia* can impact average activity levels of flies over a 3 hour period (Hague et al. 2021), our measurements of activity here over short 20-sec bouts were generally very close to zero for the majority of flies assayed (Supplemental Figure S3).

### Phylogenetic analysis

We generated *Wolbachia* and *Drosophila* phylogenies to use in the phylogenetic comparative analyses described below. The *Wolbachia* phylogeny was generated in our previous analysis of host activity (Hague et al. 2021). The Bayesian phylogram of the A-and B-group *Wolbachia* was generated using 211 single-copy genes of identical length in all *Wolbachia* genomes, spanning a total of 178,569 bp (Figure 1). To generate a *Drosophila* host phylogeny (Figure 2), publicly available *Drosophila* sequences were obtained for the three *w*Mel-*D. melanogaster* genotypes used in our experiments (Hague, Caldwell, et al. 2020; Hague et al. 2021), *D. simulans* (Hu et al., 2013), *D. mauritiana* (Meany et al. 2019), *D. sechellia* (Schrider et al. 2018), *D. yakuba*, *D. santomea*, *D. teissieri* (Cooper et al. 2019), *D. suzukii* (Turelli et al. 2018), and *D. auraria* (Conner et al. 2021). The host phylogeny was generated using the same 20 nuclear genes implemented in Turelli *et al*. (2018): *aconitase, aldolase, bicoid, ebony, enolase, esc, g6pdh, glyp, glys, ninaE, pepck, pgi, pgm, pic, ptc, tpi, transaldolase, white, wingless,* and *yellow*. We used BLAST with the *D. melanogaster* coding sequences to extract orthologs from the genomes of each host species. Sequences were then aligned with MAFFT 7. Finally, we used RevBayes and the GTR + Γ + I model partitioned by codon position and gene to accommodate potential variation in the substitution process among genes, as described in Turelli et al. (2018).

### Analysis of body mass

We tested first for interspecific variation in body mass in a comparative framework, absent potential effects from *Wolbachia*. Body masses (mg) from *Wolbachia*-negative females and males of all genotypes were used in a linear model to test for effects of host genotype and sex on body mass with the “lm” function in R (R Core Team 2023). The significance of each effect was evaluated using *F* tests and type III sum of squares using the “Anova” function in the *car* R package (Fox and Weisberg 2019). We also used a phylogenetically aware model and the *Drosophila* phylogeny to test for genotype-and sex-differences in body mass while accounting for phylogenetic relatedness. Here, an extended phylogenetic generalized least squares (E-PGLS) regression was used with the “extended.pgls” function in the *geomorph* package (Collyer and Adams 2018; Baken et al. 2021; Collyer and Adams 2024; Adams et al. 2025). E-PGLS can accommodate more than one individual per species in a PGLS analysis (as opposed to using species means), which facilitates comparisons of within-species variation in body mass (i.e., females vs. males) while accounting for phylogenetic non-independence (Adams and Collyer 2024; Glynne and Adams 2026). Again, host genotype and sex were included as explanatory variables and the “anova” function was used to test for significance.

The body masses from *Wolbachia*-negative flies were also used to test for phylogenetic signal, the degree to which body mass variation is predicted by phylogenetic relationships. Mean body mass values for females and males of each genotype were used to test for phylogenetic signal on the *Drosophila* phylogeny using Pagel’s lambda (λ). A Pagel’s λ value of 0 coincides with character evolution that is independent of phylogenetic relationships, whereas λ = 1 is consistent with a Brownian motion model of trait evolution on the phylogeny. The “phylosig” function in the *phytools* package (Revell 2024) was used to compare fitted λ values to a model assuming no phylogenetic signal (λ = 0) using a likelihood ratio test. In this analysis, we used mean body mass estimates from *Wolbachia*-negative flies from the *w*Ri-*D. simulans* genotype for the *D. simulans* tip on the *Drosophila* phylogeny; however, the means from the *w*Ha or *w*Ha-*w*No *D. simulans* genotypes did not meaningfully alter the results.

To evaluate uncertainty surrounding estimates of λ, 95% confidence intervals were generated using 1,000 bootstrap replicates in the *pmc* package (Boettiger et al., 2012). When applicable, we conducted additional analyses to evaluate whether the number of taxa in the *Drosophila* phylogeny (*N* = 11 taxa) limited our ability to detect significant departures from λ = 0 (Hague, Caldwell, et al. 2020; Hague et al. 2021; Radousky et al. 2023). Small phylogenies are likely to generate near-zero values simply by chance, not necessarily because the phylogeny is unimportant for trait evolution (Boettiger et al. 2012). To evaluate whether larger phylogenies increased the accuracy of estimation, trees were simulated using an increasing number of taxa (*N* = 25, 50) and empirical λ estimates with the “sim.bdtree” and “sim.char” functions in the *geiger* package (Harmon et al. 2008). Confidence intervals were then re-estimated using the larger simulated trees.

Next, we incorporated *Wolbachia*-positive flies into focused analyses of each genotype to evaluate whether *Wolbachia* significantly alter female and/or male body mass within a given *Wolbachia*-host system. Here, the “lm” function was used to fit separate genotype-specific linear models for each *Wolbachia*-host system with *Wolbachia* infection status, sex, and a *Wolbachia*-by-sex interaction effect as explanatory variables. The significance of each effect was evaluated using *F* tests and type III sum of squares using the “Anova” function in the *car* R package (Fox and Weisberg 2019).

Finally, we used the *Wolbachia* phylogeny (Figure 1) to test if *Wolbachia* effects on host body mass show phylogenetic signal, addressing whether closely related *Wolbachia* strains tend to have similar effects on host body mass. Here, *Wolbachia* effects on host body mass were treated as a binary variable: each *Wolbachia* strain was scored based on whether it significantly affected host body mass in the genotype-specific linear models described above. We then tested for phylogenetic signal of the binary trait using the *D* statistic (Fritz and Purvis 2010), implemented in the *caper* package (Orme et al. 2018). The *D* statistic tests for significant departures from a Brownian expectation of phylogenetic signal (*D* = 0) and a model of phylogenetic randomness (*D* = 1).

### Analysis of metabolic rate

We began the analysis of metabolic rate using the *Wolbachia*-negative flies to test whether mass-adjusted SMR varies interspecifically, absent potential effects from *Wolbachia*. To adjust for mass, body mass was included as an explanatory variable in the linear mixed model (LMM) analyses of SMR described below. Log-transformed SMR (nmol min^-1^) values from the females and males of all genotypes were used in a LMM that included host genotype, sex, log-transformed body mass, and a sex-by-mass interaction effect as fixed effects, in addition to the following nuisance variables that could plausibly affect metabolic rate: average activity (ADS), start time of the SMR recording, average flow rate in the chamber (ml min^-1^), average water vapor (ppt), average temperature (°C), and average light intensity (lux). The ID of each fly was included as a random effect to account for the fact that SMR of each fly was measured multiple times. The LMM was run using the “lmer” function in the *lme4* package (Bates et al. 2015) and significance of the fixed effects was assessed using *F* tests and type III sum of squares with a Kenward-Roger approximation for degrees of freedom using the “Anova” function and the *pbkrtest* and *car* R packages (Halekoh and Højsgaard 2014; Fox and Weisberg 2019). We also used a phylogenetically aware model to test for genotype-and sex-differences in mass-adjusted metabolic rate while accounting for phylogenetic relatedness among *Drosophila* species using the “extended.pgls” function. For the E-PGLS, only the first SMR recording from each fly was included, because E-PGLS does not accommodate mixed models.

We also tested whether mass-adjusted SMR shows evidence of phylogenetic signal using the *Drosophila* phylogeny. Least-square (LS) mean mass-adjusted SMR values for *Wolbachia*-negative females and males were extracted from the genotype-specific LMMs described below (to account for the effects of body mass and nuisance variables on SMR) and used as mean trait values in tests for phylogenetic signal using Pagel’s λ. LS means were extracted using the “emmeans” function in the *emmeans* package (Lenth 2025). For the *D. simulans* tip on the *Drosophila* phylogeny, LS mean values from the *w*Ri-*D. simulans* genotype were used; however, LS means from the *w*Ha or *w*Ha-*w*No genotypes did not meaningfully alter the results.

Next, we incorporated *Wolbachia*-positive flies into focused analyses of each genotype to test whether *Wolbachia* significantly alter female and/or male metabolic rate within a given *Wolbachia*-host system. Here, LMMs were fit for each *Wolbachia*-host system that included the following fixed effects: *Wolbachia* infection status, sex, and a *Wolbachia*-by-sex interaction effect, in addition to the nuisance variables log-body mass, activity, start time, flow rate, water vapor, temperature, and light intensity. The ID of each fly was included as a random effect. *F* tests and type III sum of squares with a Kenward-Roger approximation for degrees of freedom were used to test the significant of fixed effects.

In the above-mentioned analysis of body mass, *Wolbachia* significantly altered body mass in the majority of *Wolbachia*-host systems (see Results, Figure 3). Metabolic rates usually scale with body mass, which we confirmed with interspecific and genotype-specific analyses of mass scaling relationships (see detailed analysis in Supplemental Materials). Thus, it is plausible that *Wolbachia* effects on body mass could indirectly alter SMR (e.g., *Wolbachia* increase host body mass, which increases whole-organism SMR as a consequence). We sought to distinguish between (1) mass-dependent effects of *Wolbachia* that alter whole-organism SMR and (2) mass-independent effects of *Wolbachia* that alter mass-adjusted SMR. Therefore, each of the genotype-specific LMMs was run excluding and then including body mass as an explanatory variable to test whether *Wolbachia* alter host metabolic rates before and after accounting for body mass.

**Figure 3.**
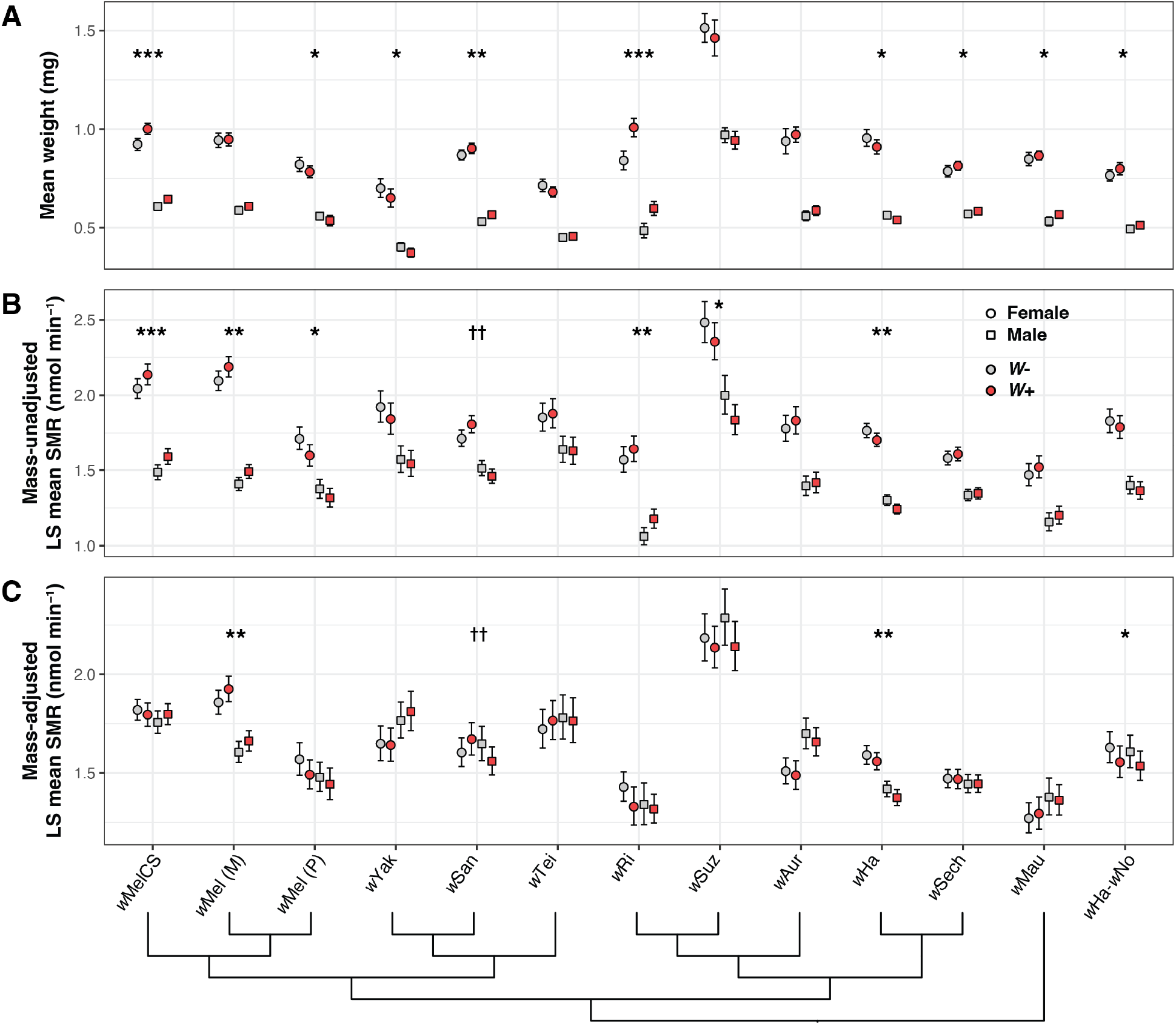
Pervasive, variable effects of *Wolbachia* on host body mass and metabolic rate. **(A)** Mean body mass (± 95% CIs) of *Wolbachia*-negative (*W*-) and -positive (*W*+) flies for each sex of each *Wolbachia*-*Drosophila* host system. **(B)** Mass-dependent effects of *Wolbachia* on SMR. LS mean mass-unadjusted SMR (± 95% CIs) of *Wolbachia*-negative and -positive flies for each sex of each genotype. **(C)** Mass-independent effects on SMR. LS mean mass-adjusted SMR (± 95% CIs) of *Wolbachia*-negative and -positive flies for each sex of each genotype. Below, the cladogram depicts evolutionary relationships among *Wolbachia* variants. Data for the co-infected *w*Ha-*w*No-*D. simulans* genotype are shown on the far right. Genotypes with a significant main effect of *Wolbachia* (*) or a *Wolbachia*-by-sex interaction (†) are marked to indicate significance (* = *P* < 0.05; ** = *P* < 0.01; *** = *P* < 0.001).

Finally, we used the *Wolbachia* phylogeny to test whether *Wolbachia* effects on metabolic rate show evidence of phylogenetic signal using the *D* statistic. As with body mass, *Wolbachia* effects on host SMR were treated as a binary variable: each *Wolbachia* strain was scored based on whether or not it had a significant effect on host SMR in the genotype-specific LMMs described above. We tested for evidence of phylogenetic signal for mass-dependent and -independent effects on SMR.

## RESULTS

### Body mass

The body masses of all *Wolbachia*-negative flies were first used to test whether *Drosophila* body mass varied interspecifically (Figure 2). In a linear model, *Drosophila* genotype (*F*_12,1530_ = 146.41, *P* < 0.001) and sex (*F*_1,1530_ = 2419, *P* < 0.001) significantly affected body mass. The sex effect reflected that females are generally larger than males (Figure 2). We also performed E-PGLS to test for variation in body mass while accounting for phylogenetic relationships among *Drosophila* species assuming a Brownian motion model of evolution. *Drosophila* genotype (*F*_12,1530_ = 10.16, *P* = 0.001) and sex (*F*_1,1530_ = 2683.12, *P* = 0.001) still had significant effects on body mass after accounting for phylogeny.

We also used mean body masses from *Wolbachia*-negative flies of each genotype to quantify the strength of phylogenetic signal on the *Drosophila* phylogeny using Pagel’s λ (Figure 2). We detected strong evidence of phylogenetic signal for both females (λ = 0.977, *P* = 0.005) and males (λ = 0.997, *P* = 0.004). This was true regardless of what genotype was included for the mean body mass estimates for the *D. simulans* tip on the tree (i.e., *Wolbachia*-negative *D. simulans* from the *w*Ri, *w*Ha, or *w*Ha-*w*No genotypes). Consistent with recent work, these results suggest that interspecific variation in *Drosophila* body mass for both sexes is strongly associated with phylogenetic relationships, but that significant variation still exists after accounting for phylogeny (Rader et al. 2026).

Next, *Wolbachia*-positive flies were incorporated into focused analyses of each *Wolbachia*-host system to evaluate how sex, *Wolbachia*, and sex-by-*Wolbachia* interactions impact host body mass using linear models (Figure 3, Supplemental Table S2). Sex had a large, significant effect on body mass in every model, again reflecting sexual size dimorphism. *Wolbachia* had a significant main effect on body mass for nine of the 13 *Wolbachia*-host systems. *Wolbachia* had a significant effect for the *w*MelCS-*D. melanogaster* genotype (F_1,456_ = 22.944, *P* < 0.001), such that *Wolbachia*-positive flies had greater body mass than -negative flies (Figure 3). This was also true for the *w*San-*D. santomea* (*F*_1,204_ = 10.607, *P* = 0.001), *w*Ri-*D. simulans* (*F*_1,125_ = 46.428, *P* < 0.001), *w*Sh-*D. sechellia* (*F*_1,206_ = 4.222, *P* = 0.041), *w*Mau-*D. mauritiana* (*F*_1,142_ = 4.652, *P* = 0.033), and co-infected *w*Ha-*w*No-*D. simulans* genotypes (*F*_1,234_ = 5.29, *P* = 0.022). We found the opposite pattern for three other genotypes, Panama *w*Mel-*D. melanogaster* (*F*_1,157_ = 4.546, *P* = 0.035), *w*Yak-*D. yakuba* (*F*_1,160_ = 4.518, *P* = 0.035), and *w*Ha-*D. simulans* (*F*_1,350_ = 5.026, *P* = 0.026), where *Wolbachia*-positive flies generally had a smaller body mass than -negative flies. There was no evidence for sex-by-*Wolbachia* interaction effects on body mass.

Finally, we used the *Wolbachia* phylogeny (Figure 1) to test for phylogenetic signal using the *D* statistic, assessing whether closely related *Wolbachia* tend to have similar effects on host body mass. The *D* value of 1.454 indicated overdisperson (*D* > 1) of effects on body mass across the tree, and a model of Brownian phylogenetic structure could be rejected (*D* = 0, *P* = 0.001). In contrast, a model of no phylogenetic structure could not be rejected (*D* = 1, *P* = 0.76), implying that *Wolbachia* effects on body mass do not show phylogenetic signal on the *Wolbachia* phylogeny.

### Metabolic rate

The estimates of SMR from all *Wolbachia*-negative flies were first used to test whether *Drosophila* metabolic rates vary interspecifically (Figure 2). The fitted LMM showed a strong effect of *Drosophila* genotype (*F*_12,1577.2 =_ 18.515, *P* < 0.001), sex (*F*_1,1530_ = 17.979, *P* < 0.001), body mass (*F*_1,1502.9_ = 371.467, *P* < 0.001), and a sex-by-mass interaction effect (*F*_1,1534_ = 371.467, *P* < 0.001). In addition, the following nuisance variables were also significant: temperature (*F*_1,2780_ = 136.796, *P* < 0.001), start time (*F*^1,1597.1^ = 1117.129, *P* < 0.001), flow rate (*F*_1,1603.1_ = 110.169, *P* < 0.001), and activity (*F*_1,2231.1_ = 418.828, *P* < 0.001). See Supplemental Table S3 for more information about nuisance variable effects. The E-PGLS incorporating *Drosophila* phylogenetic relationships indicated that *Drosophila* genotype no longer had a significant effect on SMR after accounting for phylogeny (*F*_12,1522_ = 1.492, *P* = 0.105). In contrast, sex (*F*_1,1522_ = 13.408, *P* = 0.001), body mass (*F*_1,1522_ = 198.149, *P* = 0.001), and the sex-by-mass interaction effect (*F*_12,1522_ = 4.87, *P* = 0.029) were still significant after accounting for phylogeny. Additionally, temperature (*F*_12,1522_ = 58.564, *P* = 0.001), start time (*F*_12,1522_ = 209.903, *P* = 0.001), flow rate (*F*_12,1522_ = 400.586, *P* = 0.001), and activity (*F*_12,1522_ = 138.4312, *P* = 0.001) were significant in the phylogenetically informed model.

We also used mean estimates of metabolic rate from *Wolbachia*-negative flies to characterize the strength of phylogenetic signal on the *Drosophila* phylogeny. Least-square (LS) mean mass-adjusted SMR values were extracted from the genotype-specific LMMs (described below) to test for phylogenetic signal using Pagel’s λ (Figure 2). LS mean mass-adjusted SMR values from females likely exhibit phylogenetic signal (λ = 0.926, *P* = 0.174), though the estimate was not significantly different from λ = 0. A power analysis indicated that the number of taxa in the *Drosophila* tree (*N* = 11) likely limited our ability to detect departures from λ = 0 (see Supplemental Figure S4). For males, there was strong, significant phylogenetic signal (λ = 0.947, *P* = 0.026). These results were generally similar regardless of the genotype we used for LS mean mass-adjusted SMR estimates for the *D. simulans* tip on the tree. Together, the comparative results from *Wolbachia*-negative flies indicate that much of the interspecific variation in mass-adjusted SMR can be explained by phylogenetic relationships, particularly for male flies.

Next, *Wolbachia*-positive flies were incorporated into focused analyses of each *Wolbachia*-host genotype to evaluate how sex, *Wolbachia*, and sex-by-*Wolbachia* interactions impact SMR using LMMs (Figure 3, Supplemental Tables S4, S5). Here, we compared LMM results before and after including body mass as an explanatory variable in order to distinguish *Wolbachia* effects on whole-organism SMR vs. mass-adjusted SMR. Sex had a significant main effect on SMR for all the genotypes in the mass-unadjusted models, reflecting that females generally have a higher whole-organism metabolic rate due to sexual size dimorphism. Mass-adjusted SMR varied by sex for only a few genotypes. Females had a greater mass-adjusted SMR than males for the Maine *w*Mel-*D. melanogaster* (*F* _1,327.8_ = 35.96, *P* < 0.001) and *w*Ha-*D. simulans* (*F* _1,346.1_ = 32.319, *P* < 0.001) genotypes, whereas females had a lower mass-adjusted SMR for the *w*Yak-*D. yakuba* (*F* _1,163.3_ = 5.229, *P* = 0.024) and *w*Aur-*D. auraria* (*F*_1,177.8_ = 10.986, *P* = 0.001) genotypes.

In a few *Wolbachia*-host systems, *Wolbachia* altered whole-organism SMR due to aforementioned *Wolbachia* effects on body mass and the mass scaling relationship with metabolic rate (see mass scaling analysis in Supplemental Materials). For example, *Wolbachia* increased SMR for the *w*MelCS-*D. melanogaster* genotype in the mass-unadjusted SMR model (*F*_1,452.4_ = 11.352, *P* < 0.001); however, the effect disappeared in the mass-adjusted model (*F*_1,449.7_ = 0.131, *P* = 0.718), implying that the increase in mass-unadjusted SMR is a result of *w*MelCS increases to host body mass (*F*_1,456_ = 22.944, *P* < 0.001). After accounting for body mass, we no longer detected a *Wolbachia* effect on SMR. This was also true for the *w*Ri-*D. simulans* genotype: *Wolbachia* increased SMR in the mass-unadjusted model (*F*_1,122.8_ = 7.553, *P* = 0.007), but not in the mass-adjusted model (*F*_1,122.4_ = 2.576, *P* = 0.111), implying that *w*Ri increases in whole-organism SMR are due to increases in body mass (*F*_1,125_ = 46.428, *P* < 0.001). For the Panama *w*Mel-*D. melanogaster* genotype, *Wolbachia* decreased SMR in the mass-unadjusted model (*F*_1,153.9_ = 6.101, *P* = 0.015), but not in the mass-adjusted model (*F*_1,152.7_ = 3.037, *P* = 0.0834), indicating that the decrease in whole-organism SMR is due to the fact that Panama *w*Mel decreases body mass (*F*_1,157_ = 4.546, *P* = 0.035). For the *w*Suz-*D. suzukii* genotype, we also found evidence that *Wolbachia* decreases SMR in the mass-unadjusted model (*F*_1,154.9_ = 5.78, *P* = 0.017); however, this effect disappeared in the mass-adjusted model (*F*_1,153.9_ = 3.341, *P* = 0.07) despite the fact that *w*Suz does not seem to affect host body mass (*F*_1,158_ = 1.243, *P* = 0.267).

For other systems, *Wolbachia* altered host metabolic rate in a mass-independent manner. For the Maine *w*Mel-*D. melanogaster* genotype, *Wolbachia* increased SMR in both the mass-unadjusted (*F*_1,332.5_ = 10.718, *P* = 0.001) and mass-adjusted models (*F*_1,329.6_ = 7.277, *P* = 0.007). We found the opposite pattern for the *w*Ha-*D. simulans* genotype, where *Wolbachia* decreased SMR in both the mass-unadjusted (*F*_1,345.9_ = 9.406, *P* = 0.002) and mass-adjusted models (*F*_1,344_ = 4.788, *P* = 0.029). For the *w*San-*D. santomea* genotype, there was a significant sex-by-*Wolbachia* interaction effect for both the mass-unadjusted (*F*_1,196_ = 7.566, *P* = 0.007) and mass-adjusted models (*F*_1,193.8_ = 9.462, *P* = 0.002), wherein *Wolbachia*-positive females had a greater metabolic rate than -negative females, but males showed the opposite pattern. For the co-infected *w*Ha-*w*No-*D. simulans* genotype, we detected a negative effect of *Wolbachia* on SMR only after accounting for body mass in the mass-adjusted model (mass-unadjusted: *F*_1,230.8_ = 1.305, *P* = 0.255; mass-adjusted: *F*_1,228.7_ = 5.796, *P* = 0.017). These genotype-specific effects on SMR generally did not correlate with the *Wolbachia* effects on host locomotor activity (measured over a 3-hour period) that we documented for the flies in this study (Supplemental Figure S5; <u>Hague</u> <u>et al. 2021)</u>.

Finally, we used the *Wolbachia* phylogeny (Figure 1) to test whether closely related *Wolbachia* strains tended to have similar effects on host metabolic rate. Using results from the mass-unadjusted LMMs, the estimate of *D* = 0.831 was high enough to reject a model of Brownian phylogenetic structure (*D* = 0, *P* = 0.024), but not a model of random phylogenetic structure (*D* = 1, *P* = 0.335). Similarly, the *D* = 1.422 value based on the mass-adjusted LMMs was high enough to reject a *D* = 0 model (*P* = 0.005), but not a *D* = 1 model (*P* = 0.773). Together, these results suggest that mass-dependent and -independent effects of *Wolbachia* on SMR do not show evidence of phylogenetic signal on the *Wolbachia* phylogeny.

## DISCUSSION

*Wolbachia* had pervasive effects on the body mass and metabolic rate of *Drosophila* hosts. Interspecific variation was largely driven by host phylogenetic relationships, but within a given system, body mass and/or SMR were almost always impacted by *Wolbachia*. In some instances, *Wolbachia* altered whole-organism metabolic rates indirectly due to effects on body mass. In others, *Wolbachia* directly altered metabolic rates through mass-independent effects. This variation from one host system to the next is consistent with complex effects arising from interactions among *Wolbachia* and host genomes, as documented for other traits, including *Wolbachia*-induced cytoplasmic incompatibility. Below, we identify underlying mechanisms that may explain mass-dependent and -independent effects on SMR and then discuss the implications for endosymbiont-host relationships.

### Mass-dependent effects

*Wolbachia* altered body mass for most hosts, including genotypes carrying A-group *Wolbachia*, the divergent B-group, and the *w*Ha-*w*No co-infection (Figure 3). In some instances, *Wolbachia* effects on body mass also altered whole-organism SMR. In most cases, we found that *Wolbachia* increased body mass, which is consistent with previous studies focused mostly on *w*Mel in *D. melanogaster*. For example, *w*Mel has been associated with increased female body mass (Kriesner et al. 2016) and, on nutrient-poor food, *w*Mel increased pupae size and adult emergence rates (Lindsey et al. 2025). Outside of *w*Mel-*D. melanogaster*, female *D. innubila* carrying *w*Inn (another A-group strain) were slightly larger and showed higher fecundity when reared on nutrient-poor food (Unckless and Jaenike 2012; Hill et al. 2022).

Such findings have led to the hypothesis that facultative *Wolbachia* are involved in nutrient provisioning (Zug and Hammerstein 2015a; Newton and Rice 2020), a phenomenon observed in obligate *Wolbachia* relationships (Foster et al. 2005; Hosokawa et al. 2010; Nikoh et al. 2014). Positive effects on host growth could arise through several routes. For instance, genes associated with the heme biosynthetic pathway are generally found in *Wolbachia* genomes (Newton and Rice 2020), and *w*Mel-positive *D. melanogaster* have increased fecundity when reared on iron-restricted or -overloaded diets (Brownlie et al. 2007; Brownlie et al. 2009). *Wolbachia* are also involved in synthesis of the B vitamins biotin and/or riboflavin in a number of host species (Hosokawa et al. 2010; Nikoh et al. 2014; Moriyama et al. 2015; Gerth and Bleidorn 2017; Preuss et al. 2026). Riboflavin is an essential cofactor for mitochondrial electron-transport-chain enzymes, and *Wolbachia*-synthesized riboflavin is linked to accelerated ovarian development and higher energy production in the rice planthopper *Laodelphax striatellus* carrying the *w*Stri strain (group B) (Bing et al. 2020; Niu et al. 2025). Finally, removal of *w*Mel from *D. melanogaster* exacerbated insulin/insulin growth factor-like signaling (IIS) mutant phenotypes, implying that *w*Mel may increase IIS in *Wolbachia*-positive flies (Ikeya et al. 2009).

Certain *Wolbachia*-host systems (Panama *w*Mel, *w*Yak, and *w*Ha) showed the opposite pattern, in which *Wolbachia* decreased body mass. These results highlight the complexity of our findings. Endosymbionts may increase growth under certain conditions, but they also likely consume host nutrients (Clavé et al. 2022). *Wolbachia* titer can change in response to host diet, implying that *Wolbachia* rely on nutrients from host cells (Ponton et al. 2015; Serbus et al. 2015). Increased food intake has been documented for *w*Mel-positive males and, after 24 hours of fasting, males had lower body mass than did *Wolbachia*-negative flies, suggesting that *w*Mel requires increased host energy expenditure (Zhang et al. 2021). Genomic and metabolomic analyses indicate that *w*Mel interacts with a wide range of host metabolic pathways in *D. melanogaster*, including aspects of amino acid, carbohydrate, and lipid metabolism (Wu et al. 2004; Jiménez et al. 2019; Carneiro Dutra et al. 2020; Currin-Ross et al. 2021; Zhang et al. 2021; Lindsey et al. 2025). *w*Mel may also depress IIS pathways (contrary to the results described above), which could help explain decreases in body mass (Haqshenas et al. 2019; Currin-Ross et al. 2021). It is also plausible that if *Wolbachia* impact the composition of host microbiota, this could have negative effects on body mass (Simhadri et al. 2017; Carneiro Dutra et al. 2020). Together, this work suggests that *Wolbachia* have widespread effects on body mass that likely depend on the context, including the physiological state of the host.

### Mass-independent effects

We also detected mass-independent effects of *Wolbachia* on SMR. *w*Mel increased mass-adjusted SMR for the Maine *D. melanogaster* genotype, whereas *w*Ha and the *w*Ha-*w*No coinfection depressed it in *D. simulans*. For the *w*San-*D. santomea* genotype, *Wolbachia* increased mass-adjusted SMR in females but decreased it in males. We hypothesize that these complicated mass-independent effects result from interactions between *Wolbachia* and mitochondria. *Wolbachia* are maternally co-transmitted with mitochondria (Henry and Newton 2018; Russell et al. 2019; Porter and Sullivan 2023; Radousky et al. 2023; Hague et al. 2024), and multiple lines of evidence imply that *Wolbachia* interact with mitochondria. For example, RNAi knockdown of multiple mitochondrial genes altered *Wolbachia* titer in *Drosophila* cell lines and developing oocytes (White et al. 2017), and *w*Mel altered expression of host genes related to mitochondrial metabolism in *D. melanogaster* larvae (Lindsey et al. 2025). *w*MelCS also altered expression of glutamate dehydrogenase, a key enzyme in the TCA cycle, as well as ATP levels in *D. melanogaster* (Carneiro Dutra et al. 2020).

*Wolbachia* can also affect levels of reactive oxygen species (ROS) and the host oxidative environment, which may reflect *Wolbachia* interactions with the host immune system. These effects have been documented for *w*MelCS in *D. melanogaster* and *w*Ri in *D. simulans* (Brennan et al. 2008; Brennan et al. 2012; Pan et al. 2012; Wong et al. 2015; Zug and Hammerstein 2015b; Monnin et al. 2016). Similarly, *Wolbachia* interactions with host iron metabolism could plausibly affect ROS levels and mitochondrial function (Kremer et al. 2009; Gill et al. 2014; Currin-Ross et al. 2021).

### Complexity of host effects

We found that *Wolbachia* have widespread—but variable—effects on key physiological traits. Comparative studies of diverse *Drosophila* species have documented similar patterns of complex variation in the strength of *Wolbachia*-induced cytoplasmic incompatibility (Shropshire et al. 2022; Bagchi et al. 2026), maternal transmission fidelity (Radousky et al. 2023; Hague et al. 2024), and *Wolbachia* effects on host behavior (Hague, Caldwell, et al. 2020; Hague et al. 2021). We hypothesize that *Wolbachia* interactions with the host genomes underlie complex variation in host effects. For instance, the Maine and Panama *w*Mel variants are very similar (zero third-position pairwise differences across 740 single copy genes, 757,926 total bp), yet differed in their effects on body mass and metabolic rate in their natural *D. melanogaster* backgrounds. Host backgrounds are known to modulate *Wolbachia* effects on host fecundity (Olsen et al. 2001), the strength of cytoplasmic incompatibility (Reynolds and Hoffmann 2002; Cooper et al. 2017), and maternal transmission rates (Hague et al. 2022). *Wolbachia* titers can also vary substantially across host backgrounds (Veneti et al. 2004; Mouton et al. 2007; White et al. 2017; Funkhouser-Jones et al. 2018; Grobler et al. 2018; Hague et al. 2022), and titer differences in somatic tissues from one host system to the next could have varying effects on host physiology. Extending our analyses to host genotypes with reciprocally introgressed nuclear and cytoplasmic genomes (e.g., Hague et al. 2020; Hague et al. 2022) is required to distinguish nuclear and cytoplasmic contributions to *Wolbachia* effects on body mass and metabolic rate.

Recent host switching may also help explain the variable patterns we observed. *Wolbachia*, including variants in our study, are known to rapidly switch between divergent host species (O’Neill et al. 1992; Raychoudhury et al. 2009; Ahmed et al. 2015; Gerth and Bleidorn 2017; Turelli et al. 2018; Cooper et al. 2019; Shropshire et al. 2026). For instance, *w*Mel and related “*w*Mel-like” variants (Figure 1) diverged only 10^5^–10^6^ years ago, yet occur in hosts diverged 350 MYA spanning holometabolous insects (Shropshire et al. 2026). *w*Ri and related variants—including *w*Suz and *w*Aur—have spread even more rapidly across divergent *Drosophila* (Turelli et al. 2018), and the *w*Ha, *w*Sech, *w*No, and *w*Mau strains found within the *D. simulans* host clade (Figure 2) have histories inconsistent with cladogenic acquisition (Ballard 2000; Meany et al. 2019). Host switching generates novel genomic combinations that could plausibly affect mitochondrial function, energy production, and interactions with the nuclear genome. Cytonuclear interactions between mitochondria and the nuclear background are known to affect metabolism (Arnqvist et al. 2010; Montooth et al. 2010; Mossman et al. 2016), and future work should examinine how *Wolbachia* impact these interactions. Somatic *Wolbachia* injections can be used to decouple the association between *Wolbachia* and host mitochondria in the cytoplasm (Frydman et al. 2006; Frydman 2007), which will be necessary for dissecting effects on host metabolism. Tripartite interactions among *Wolbachia*, mitochondria, and the nuclear genome are also likely to influence the physiology of transinfected hosts. Understanding these effects has the potential to improve the application efficacy of variants like *w*Mel that block dengue and other arboviruses in mosquitoes (Hoffmann et al. 2011; Walker et al. 2011; Ross et al. 2019).

## Conclusions

Patterns of *Wolbachia* spread in natural host populations indicate that *Wolbachia* confer a fitness benefit to hosts, yet little is known about such benefits in nature (Weeks et al. 2007; Hamm et al. 2014; Cooper et al. 2017; Meany et al. 2019; Ross et al. 2019; Hague, Mavengere, et al. 2020; Cogni et al. 2021; Hague et al. 2022; Hoffmann and Cooper 2024; Graham et al. 2025). *Wolbachia* effects on body mass and metabolic rate, two central host traits, are likely to have cascading effects on aspects of host fitness, including growth and survival (Chown and Gaston 2010; Burton et al. 2011; White and Kearney 2013). Body mass effects in particular are likely to impact other aspects of host biology, because so many physiological and ecological traits are correlated with body size (Brown et al. 2004; Chown and Gaston 2010; White and Kearney 2011). *Wolbachia* effects on SMR—a measure of the cost of self-maintenance—are also likely to have wide-ranging implications for hosts. Ultimately, these metabolic effects contribute to the overall impact of *Wolbachia* on host fitness, which determines endosymbiont spread and persistence in host populations (Hoffmann et al. 1990; Kriesner et al. 2016; Hoffmann and Cooper 2024; Graham et al. 2025).

Our findings indicate that future studies involving arthropod body mass, metabolism, and related life-history traits should consider the presence of facultative endosymbionts. A growing body of work shows that facultative *Wolbachia* alter a range of important host traits with fitness implications, including fecundity (Hoffmann et al. 1990; Olsen et al. 2001; Fry et al. 2004; Weeks et al. 2007), nutrition (Brownlie et al. 2009; Hosokawa et al. 2010; Nikoh et al. 2014; Newton and Rice 2020; Lindsey et al. 2025), immune function (Hedges et al. 2008; Teixeira et al. 2008; Osborne et al. 2009; Martinez et al. 2014; Cogni et al. 2021; Bruner-Montero and Jiggins 2023; Porter and Sullivan 2023), temperature preference (Arnold et al. 2019; Truitt et al. 2019; Hague, Caldwell, et al. 2020), and activity levels and sleep (Vale and Jardine 2015; Bi et al. 2018; Morioka et al. 2018; Hague et al. 2021). Elucidating how *Wolbachia* affect host traits like body mass and metabolic rate is critical for understanding host-endosymbiont evolution in natural systems and for optimizing endosymbiont-based applications in transinfected vector and pest systems (Hoffmann and Cooper 2025).

## DATA AVAILABILITY

All data and code are available at https://github.com/jmgraham30/Metabolism_final_R_scripts.

## AUTHOR CONTRIBUTIONS

JMG: data curation (lead), formal analysis (equal), visualization (equal), writing – review and editing (supporting). HAW: conceptualization (equal), methodology (lead), writing – review and editing (supporting). BSC: conceptualization (equal), project administration (equal), funding acquisition (lead), supervision (lead), writing – review and editing (supporting). MTJH: project administration (equal), data curation (supporting), formal analysis (equal), visualization (equal), supervision (supporting), writing – original draft (lead), writing – review and editing (lead).

## AKNOWEDGEMENTS

We thank Tim Wheeler and Isaac Humble for laboratory assistance and Will Conner for help with bioinformatic analyses. We are also grateful to Romain Boisseau for guidance with flowthrough respirometry.

