## Supplemental Materials for "Pervasive effects of *Wolbachia* on host body mass and metabolic rate"

#### Analysis of mass scaling relationships

*Methods*

The relationship between metabolic rate ($SMR$) and body mass ($M$) is typically expressed as a power function $SMR = aM^b$, such that $\log(SMR) = \log(a) + b\log(M)$ and $b$ represents the mass scaling exponent ($b$ = 1 represents isometry). The *Wolbachia*-negative flies from all genotypes were first used to estimate an interspecific scaling exponent. The "emtrends" function in the *emmeans* package (Lenth 2025) was used to extract sex-specific slopes and confidence intervals from the interspecific LMM analysis of mass-adjusted SMR, which describe the scaling relationship between log-SMR and log-mass for each sex.

We also conducted focused analyses of each genotype to examine whether *Wolbachia* alter the relationship between metabolic rate and body mass for either sex within a given *Wolbachia*-host system. For each genotype, the "lmer" function was used to fit a LMM with log-SMR as the dependent variable and a three-way sex-by-infection-by-log-body mass interaction as a fixed effect (in addition to the previously mentioned nuisance variables and fly ID as a random effect). This allowed for the extraction of slopes for *Wolbachia*-positive and negative flies of each sex that describe the relationship between log-SMR and log-mass. When considering intraspecific data, metabolic rate typically varies more than body mass, and a narrow range of body mass values can make it difficult to estimate mass scaling exponents (Somjee et al. 2021). To quantify body mass variation, we also calculated the body mass size range, which represents the order of magnitude difference between the largest and smallest individuals measured (log(*max*) - log(*min*)) (Chown et al. 2007).

*Results*

The *Wolbachia*-negative flies were first used to examine interspecific scaling relationships between body mass and metabolic rates in *Drosophila* species. The interspecific LMM analysis showed that SMR scaled hypometrically ($\beta < 1$) across species for both females ($\beta$ = 0.533 [0.479, 0.587]) and males ($\beta$ = 0.404 [0.340, 0.467]) (Supplemental Figure S6). These results indicate that, across species, larger flies have higher metabolic rates than smaller flies, but metabolic rates do not change in proportion to the difference in body mass. This was more true for males, because they had a smaller mass scaling exponent.

In focused analyses of each genotype, scaling exponents generally did not vary by sex or *Wolbachia* status based on overlapping confidence intervals (Supplemental Figures S5, S7, Table S6). We found that some genotypes with narrow body mass size ranges below ~0.3 made it difficult to accurately estimate scaling exponents that were significantly different from a slope of zero. Narrow size ranges are not unexpected given that we used isogenic inbred lines. Scaling exponent confidence intervals that overlapped with zero included *Wolbachia*-negative (body mass size range = 0.252; $\beta$ = 0.078 [-0.202, 0.358]) and -positive male *D. santomea* (range = 0.192; $\beta$ = -0.037 [-0.441, 0.366]), -negative male *D. teissieri* (range = 0.332; $\beta$ = 0.128 [-0.225, 0.482]), -negative male *D. suzukii* (range = 0.221; $\beta$ = 0.243 [-0.237, 0.722]), -positive female *D. mauritiana* (range = 0.181; $\beta$ = 0.269 [-0.251, 0.788]), and -negative males from the *w*Ha-*w*No-*D. simulans* genotype (range = 0.273; $\beta$ = -0.355 [-0.725, 0.013]). The *D. simulans* scaling exponent stood out because the slope was quite negative; however, this was largely driven by a male with an unusually large mass value (0.774 mg). After removing the individual from the dataset, the slope for *Wolbachia*-negative *D. simulans* males was closer to zero (range = 0.160; $\beta$ = 0.069 [-0.399, 0.537]). Removing this large individual did not alter our prior conclusions about *Wolbachia* effects on body mass or SMR.

The *w*Aur-*D. auraria* genotype was the only species with a sex difference in scaling exponents based on non-overlapping confidence intervals. Here, *Wolbachia*-negative females had a greater slope ($\beta$ = 0.89 [0.743, 1.037]) than males ($\beta$ = 0.466 [0.251, 0.681]), indicating that female body mass and metabolic rate scale close to isometry, while males scale hypometrically. When comparing *Wolbachia*-negative and -positive flies from each genotype, we detected very little variation in scaling exponents. *Wolbachia*-positive males from the *w*Ha-*w*No-*D. simulans* genotype had a larger mass scaling exponents ($\beta$ = 0.682 [0.378, 0.986]) than -negative males ($\beta$ = -0.355 [-0.725, 0.013]); however, this was no longer the case when the one individual with a large mass value was removed from the dataset ($\beta$ = 0.069 [-0.399, 0.537]). Otherwise, there was no evidence that mass scaling exponents were altered by *Wolbachia* in either females or males.

### Supplemental Figures

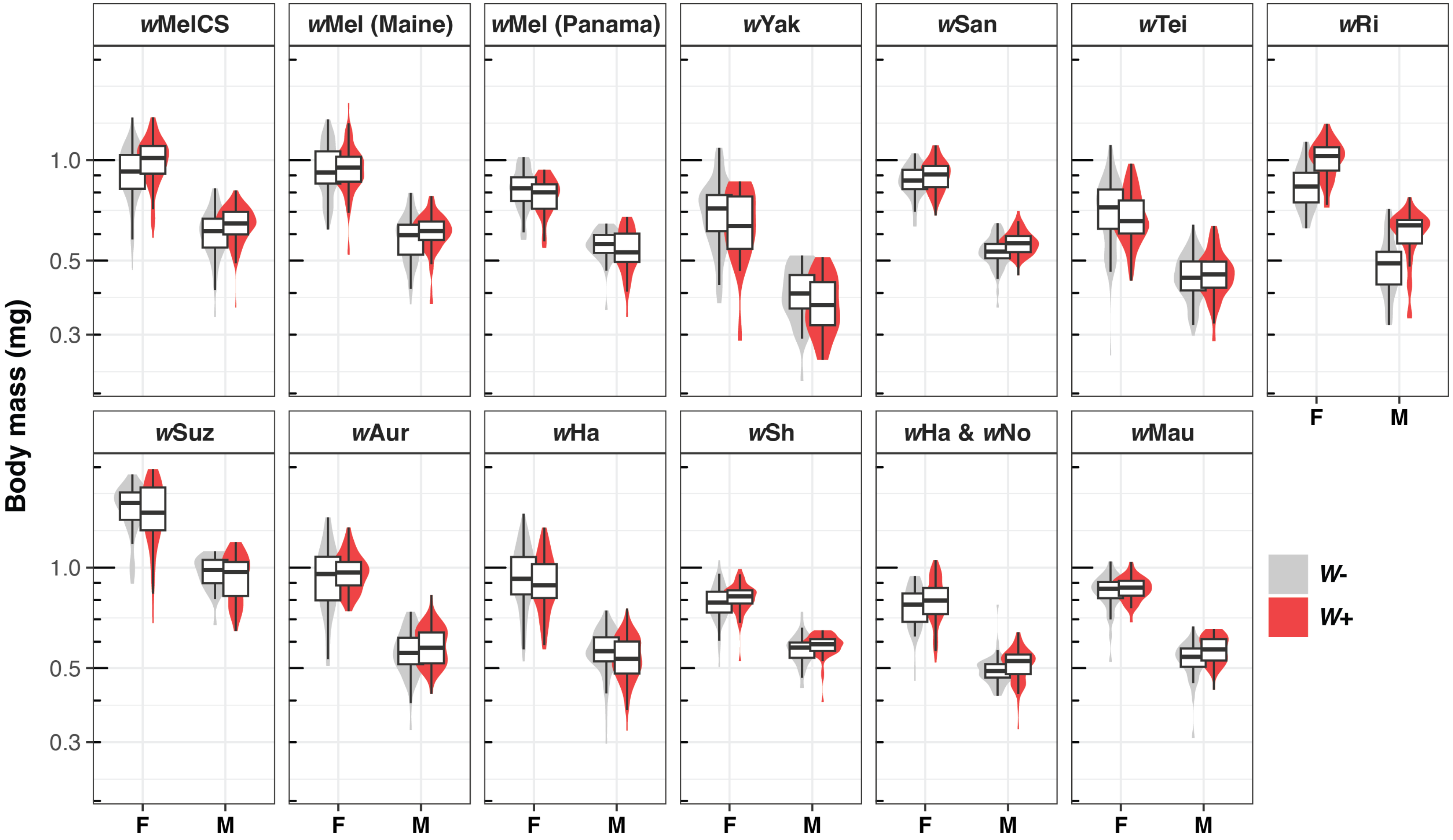


**Supplemental Figure S1.** Violin and box plots showing the distribution of *Drosophila* body masses (mg) for *Wolbachia*-negative (*W*-) and -positive (*W*+) females (F) and males (M) of each *Wolbachia-Drosophila* genotype.

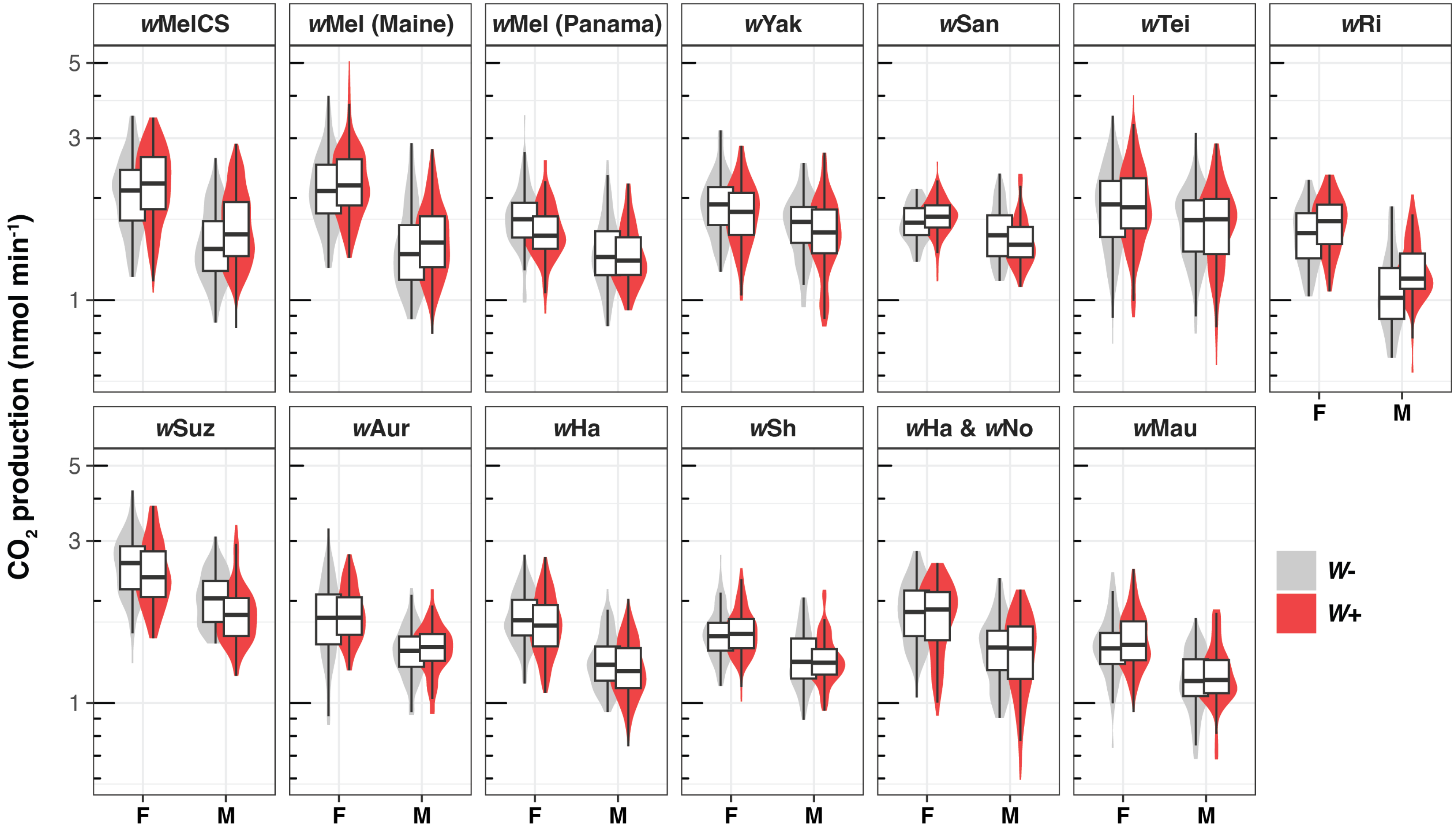


**Supplemental Figure S2.** Violin and box plots showing the distribution of *Drosophila* standard metabolic rate (SMR) measurements quantified as the lowest average 20-sec continuous bout of $CO_2$ production (nmol min$^{-1}$) for *Wolbachia*-negative (*W*-) and -positive (*W*+) females (F) and males (M) of each *Wolbachia*-*Drosophila* genotype.

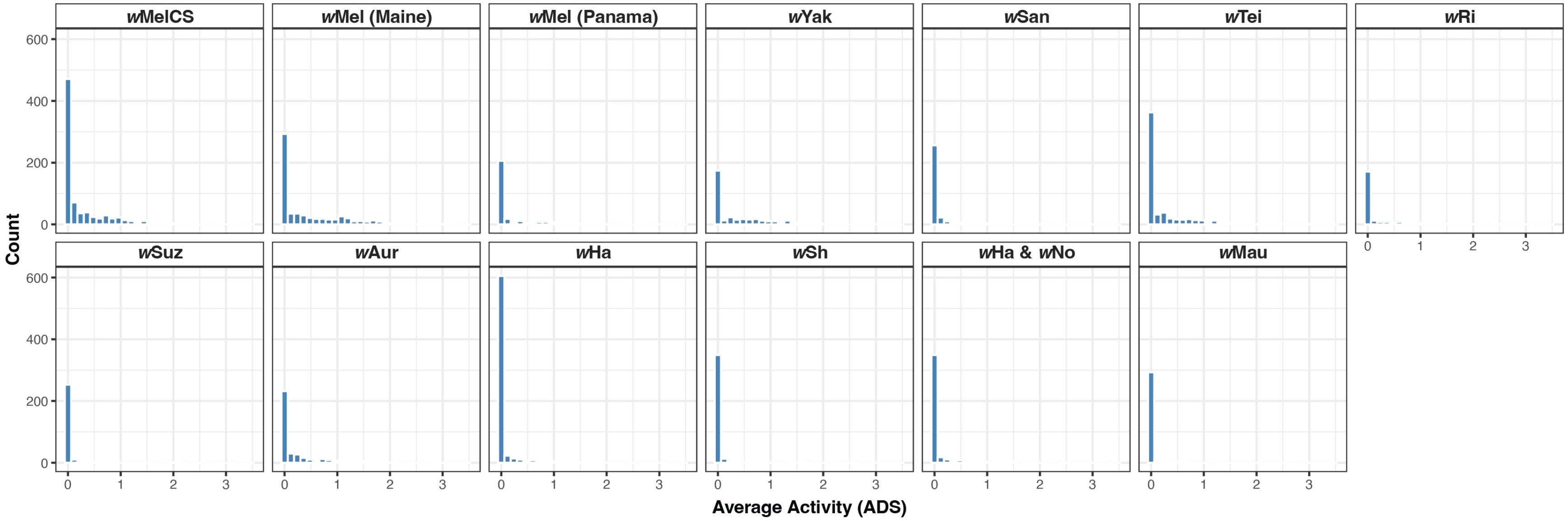


**Supplemental Figure S3.** Histograms showing the distribution of average activity measurements (ADS) for flies from each *Wolbachia-Drosophila* genotype over the 20-sec bouts when SMR was recorded.

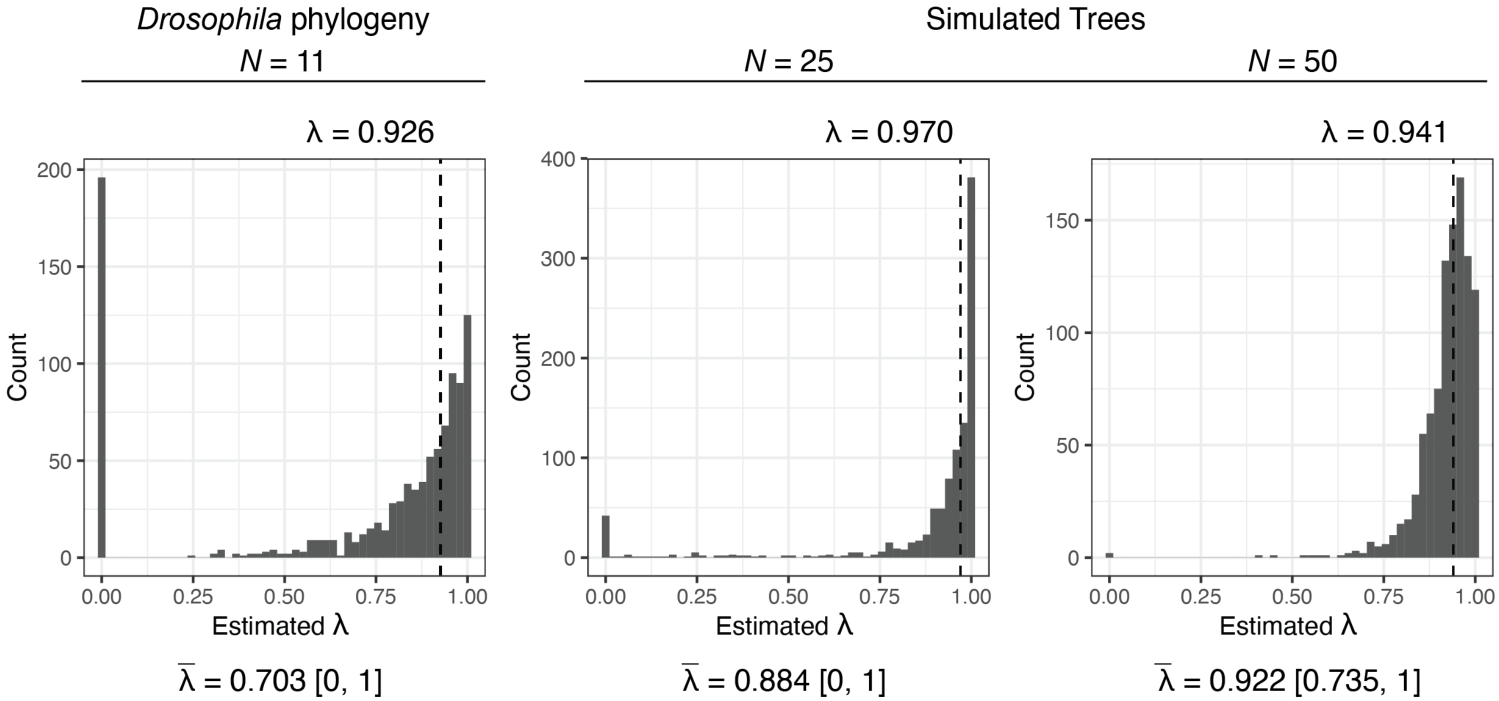


**Supplemental Figure S4.** Distribution of maximum likelihood estimates of $\lambda$ from 1,000 bootstrap replicates. The bootstrap analysis for the *Drosophila* phylogeny (Figure 2 in main text) is show to the left. To the right are simulated phylogenies with an increasing number of taxa included ($N = 25, 50$). For simulated trees, character evolution was simulated with our $\lambda$ estimate of 0.926 using the "sim.bdtree" and "sim.char" functions in the *geiger* R package (Harmon et al. 2008). For each graph, fitted $\lambda$ values for the original phylogeny are shown above with a vertical dashed line. Note that fitted $\lambda$ values for the simulated phylogenies differ slightly from $\lambda = 0.926$, because "sim.char" uses a Brownian-motion model to simulate character evolution along the phylogeny. Below each graph, the mean estimate of $\lambda$ from the 1,000 replicates ($\bar{\lambda}$) is shown with associated 95% confidence intervals. The bootstrapping analyses show that smaller phylogenies ($N \leq 25$ taxa) have a large number of near-zero $\lambda$ values arising by random chance, which increases the uncertainty of parameter estimation. Small phylogenies are known to generate near-zero $\lambda$ values by chance, not necessarily because the phylogeny is unimportant for trait evolution (Boettiger et al. 2012). As the number of taxa in our analysis increases ($N = 50$), bootstrapped estimates of $\lambda$ cluster around the true $\lambda$ value fitted to the original phylogeny.

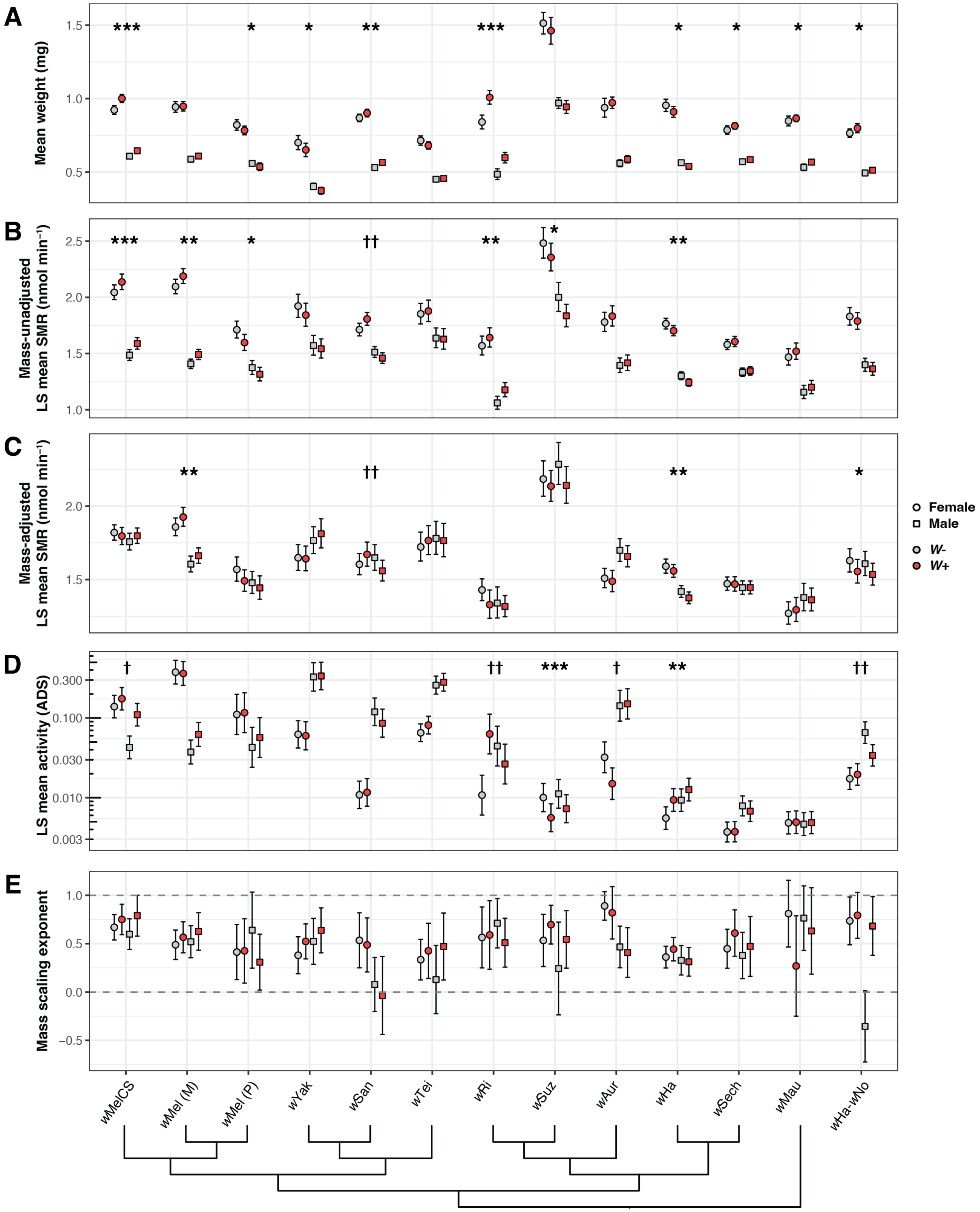


**Supplemental Figure S5.** Results from all analyses of *Wolbachia* effects on *Drosophila* host metabolism and activity. **(A)** Mean body mass (± 95% CIs) of *Wolbachia*-negative (*W*-) and -positive (*W*+) flies for each sex of each *Wolbachia-Drosophila* host system. **(B)** LS mean massunadjusted SMR (± 95% CIs) of *Wolbachia*-negative and -positive flies for each sex of each genotype. **(C)** LS mean mass-adjusted SMR (± 95% CIs) of *Wolbachia*-negative and -positive flies for each sex of each genotype. **(D)** LS mean locomotor activity (± 95% CIs) over a 3-hour period for *Wolbachia*-negative and -positive flies for each sex of each genotype. The y-axis is plotted on a log scale. Data are reproduced from Hague et al. (2021). **(E)** Mass scaling exponents (± 95% CIs) of *Wolbachia*-negative and -positive flies for each sex of each genotype. Slope values of 0 (no relationship between log-mass and log-SMR) and 1 (isometry) are marked with horizontal dashed lines. Below, the cladogram depicts evolutionary relationships among *Wolbachia* variants. Data for the co-infected *w*Ha-*w*No-*D. simulans* genotype are shown on the far right. Genotypes with a significant main effect of *Wolbachia* (*) or a *Wolbachia*-by-sex interaction (†) are marked to indicate significance (* = $P < 0.05$; ** = $P < 0.01$; *** = $P < 0.001$).

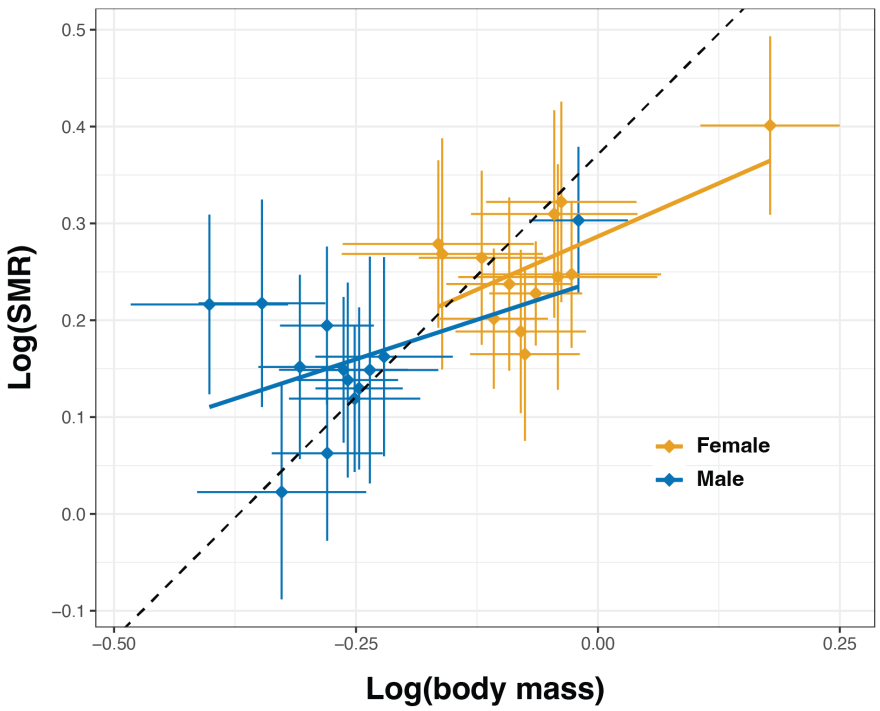


**Supplemental Figure S6.** Interspecific analysis of log-transformed body mass (mg) and log-transformed metabolic rate (nmol min$^{-1}$) values from *Wolbachia*-negative flies. Diamonds show means with ± 1 s.d. error bars for females and males of each genotype. Solid lines represent linear regression lines of log(SMR) on log(body mass) for each sex. The dashed line represents isometry ($\beta$ = 1) with an arbitrary intercept.

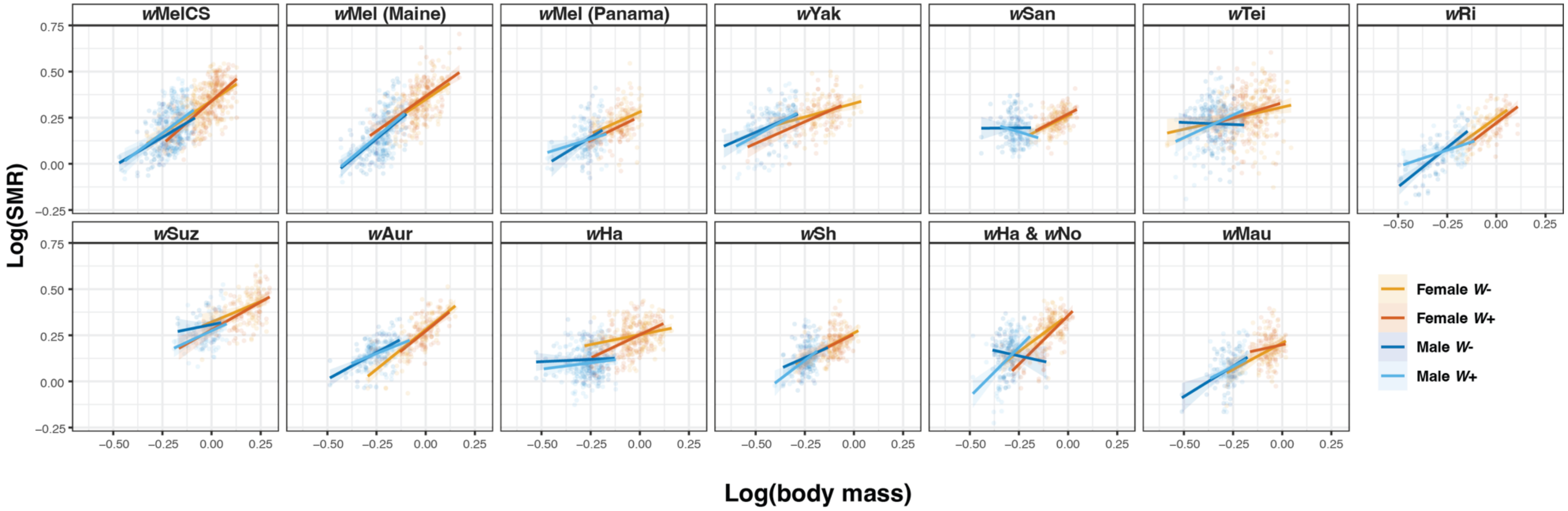


**Supplemental Figure S7.** Genotype-specific analyses of log-body mass (mg) and log-SMR (nmol min$^{-1}$) values for *Wolbachia*-negative (*W*-) and -positive (*W*+) females and males of each *Wolbachia-Drosophila* genotype. Solid lines represent linear regression lines of log(SMR) on log(body mass) for each treatment group.

**Supplemental Tables**

**Supplemental Table S1.** Isofemale lines used in analyses of *Drosophila* body mass and metabolic rate.

| **Species** | ***Wolbachia*** | **Genotype ID** | **Collection Location** |
|---|---|---|---|
| *D. melanogaster* | *w*MelCS | *CSBerk* | Canton, Ohio, USA |
| *D. melanogaster* | *w*Mel | *FFD25* | Maine, USA |
| *D. melanogaster* | *w*Mel | *PC75* | Panama City, Panama |
| *D. yakuba* | *w*Yak | *B13L5* | Bioko, Equatorial Guinea |
| *D. santomea* | *w*San | *S09L4* | São Tomé, São Tomé and Príncipe |
| *D. teissieri* | *w*Tei | *B13L11* | Bioko, Equatorial Guinea |
| *D. simulans* | *w*Ri | *Riv84* | Riverside, California, USA |
| *D. simulans* | *w*Ha | *Car5* | Hawaii, USA |
| *D. simulans* | *w*Ha & *w*No | *sim198* | New Caledonia |
| *D. suzukii* | *w*Suz | *S278* | California, USA |
| *D. auraria* | *w*Aur | *L2* | Japan |
| *D. sechellia* | *w*Sh | *PmuseumbananaI* | Praslin, Seychelles |
| *D. mauritiana* | *w*Mau | *mauR31* | Mauritius |

**Supplemental Table S2.** Results from linear models evaluating the effects of sex and *Wolbachia* on *Drosophila* body mass.

| | | **Sex** | | | ***Wolbachia*** | | | **Sex-by-*Wolbachia*** | | |
|---|---|---|---|---|---|---|---|---|---|---|
| ***Wolbachia* Genotype** | **N** | **F** | **p value** | | **F** | **p value** | | **F** | **p value** | |
| *w*MelCS | 460 | 785.891 | 0.000 | *** | 22.944 | 0.000 | *** | 2.997 | 0.084 | ns |
| *w*Mel (Maine) | 342 | 643.889 | 0.000 | *** | 0.863 | 0.354 | ns | 0.383 | 0.536 | ns |
| *w*Mel (Panama) | 161 | 330.157 | 0.000 | *** | 4.546 | 0.035 | * | 0.267 | 0.606 | ns |
| *w*Yak | 164 | 243.071 | 0.000 | *** | 4.518 | 0.035 | * | 0.315 | 0.575 | ns |
| *w*San | 208 | 1056.021 | 0.000 | *** | 10.607 | 0.001 | ** | 0.003 | 0.957 | ns |
| *w*Tei | 346 | 393.789 | 0.000 | *** | 1.283 | 0.258 | ns | 2.482 | 0.116 | ns |
| *w*Ri | 129 | 346.732 | 0.000 | *** | 46.428 | 0.000 | *** | 1.781 | 0.185 | ns |
| *w*Suz | 162 | 229.655 | 0.000 | *** | 1.243 | 0.267 | ns | 0.128 | 0.721 | ns |
| *w*Aur | 182 | 355.796 | 0.000 | *** | 2.243 | 0.136 | ns | 0.023 | 0.879 | ns |
| *w*Ha | 354 | 628.379 | 0.000 | *** | 5.026 | 0.026 | * | 0.459 | 0.499 | ns |
| *w*Sech | 210 | 470.781 | 0.000 | *** | 4.222 | 0.041 | * | 0.434 | 0.511 | ns |
| *w*Ha & *w*No | 238 | 586.269 | 0.000 | *** | 5.290 | 0.022 | * | 0.398 | 0.529 | ns |
| *w*Mau | 146 | 618.806 | 0.000 | *** | 4.652 | 0.033 | * | 0.504 | 0.479 | ns |

**Supplemental Table S3.** Regression coefficients for nuisance variables from the linear mixed model (LMM) examining interspecific variation in SMR. The explanatory variables of interest, sex and body mass, are shown at the top for reference.

| **Explanatory variable** | **Estimate** | **Std. Error** | **t value** |
|---|---|---|---|
| Sex (male) | -0.038 | 0.009 | -4.240 |
| Body mass (mg) | 0.533 | 0.028 | 19.274 |
| Temperature (°C) | 0.038 | 0.003 | 11.704 |
| Light intensity (lux) | 0.000 | 0.000 | -1.429 |
| SMR recording start time | -0.082 | 0.002 | -33.432 |
| Flow rate (ml $min^{-1}$) | -0.004 | 0.000 | -10.497 |
| Water vapor (ppt) | -0.001 | 0.000 | -1.220 |
| Fly activity (ADS) | 0.059 | 0.003 | 20.477 |

**Supplemental Table S4.** Results from LMMs evaluating the effects of sex and *Wolbachia* on SMR without adjusting for body mass.

| | | Sex | | | ***Wolbachia*** | | | Sex-by-***Wolbachia*** | | |
|---|---|---|---|---|---|---|---|---|---|---|
| ***Wolbachia* Genotype** | **N** | **F** | **p value** | | **F** | **p value** | | **F** | **p value** | |
| *w*MelCS | 460 | 344.692 | 0.000 | *** | 11.352 | 0.001 | *** | 0.468 | 0.494 | ns |
| *w*Mel (Maine) | 342 | 622.446 | 0.000 | *** | 10.718 | 0.001 | ** | 0.218 | 0.641 | ns |
| *w*Mel (Panama) | 161 | 81.010 | 0.000 | *** | 6.101 | 0.015 | * | 0.311 | 0.578 | ns |
| *w*Yak | 164 | 44.517 | 0.000 | *** | 1.184 | 0.278 | ns | 0.187 | 0.666 | ns |
| *w*San | 208 | 99.850 | 0.000 | *** | 0.298 | 0.586 | ns | 7.566 | 0.007 | ** |
| *w*Tei | 346 | 24.324 | 0.000 | *** | 0.022 | 0.883 | ns | 0.130 | 0.719 | ns |
| *w*Ri | 129 | 180.549 | 0.000 | *** | 7.553 | 0.007 | ** | 1.174 | 0.281 | ns |
| *w*Suz | 162 | 65.893 | 0.000 | *** | 5.780 | 0.017 | * | 0.337 | 0.563 | ns |
| *w*Aur | 182 | 104.829 | 0.000 | *** | 0.846 | 0.359 | ns | 0.080 | 0.778 | ns |
| *w*Ha | 354 | 523.977 | 0.000 | *** | 9.406 | 0.002 | ** | 0.152 | 0.697 | ns |
| *w*Sech | 210 | 148.565 | 0.000 | *** | 0.802 | 0.372 | ns | 0.075 | 0.784 | ns |
| *w*Ha & *w*No | 238 | 159.263 | 0.000 | *** | 1.305 | 0.255 | ns | 0.007 | 0.934 | ns |
| *w*Mau | 146 | 89.521 | 0.000 | *** | 2.071 | 0.152 | ns | 0.003 | 0.955 | ns |

| | | Temperature (C°) | | | Light (lux) | | | Start time | | | Flow rate (ml $min^{-1}$) | | | Water vapor pressure (ppt) | | | Activity (ADS) | | |
|---|---|---|---|---|---|---|---|---|---|---|---|---|---|---|---|---|---|---|---|
| ***Wolbachia* Genotype** | **N** | **F** | **p value** | | **F** | **p value** | | **F** | **p value** | | **F** | **p value** | | **F** | **p value** | | **F** | **p value** | |
| *w*MelCS | 460 | 0.245 | 0.621 | ns | 5.536 | 0.019 | * | 758.053 | 0.000 | *** | 10.561 | 0.001 | ** | 38.691 | 0.000 | *** | 105.519 | 0.000 | *** |
| *w*Mel (Maine) | 342 | 35.752 | 0.000 | *** | 0.470 | 0.493 | ns | 838.661 | 0.000 | *** | 77.444 | 0.000 | *** | 2.044 | 0.154 | ns | 48.907 | 0.000 | *** |
| *w*Mel (Panama) | 161 | 20.395 | 0.000 | *** | 0.254 | 0.615 | ns | 229.558 | 0.000 | *** | 18.561 | 0.000 | *** | 26.065 | 0.000 | *** | 86.437 | 0.000 | *** |
| *w*Yak | 164 | 8.731 | 0.003 | ** | 4.188 | 0.042 | * | 6.427 | 0.012 | * | 0.837 | 0.362 | ns | 0.042 | 0.839 | ns | 53.998 | 0.000 | *** |
| *w*San | 208 | 27.853 | 0.000 | *** | 0.843 | 0.359 | ns | 26.620 | 0.000 | *** | 0.311 | 0.578 | ns | 8.006 | 0.005 | ** | 68.068 | 0.000 | *** |
| *w*Tei | 346 | 2.304 | 0.130 | ns | 8.990 | 0.003 | ** | 20.016 | 0.000 | *** | 0.034 | 0.854 | ns | 0.393 | 0.531 | ns | 53.590 | 0.000 | *** |
| *w*Ri | 129 | 0.155 | 0.694 | ns | 0.029 | 0.866 | ns | 41.099 | 0.000 | *** | 34.419 | 0.000 | *** | 3.008 | 0.085 | ns | 54.956 | 0.000 | *** |
| *w*Suz | 162 | 8.858 | 0.003 | ** | 6.033 | 0.015 | * | 102.496 | 0.000 | *** | 0.069 | 0.793 | ns | 2.664 | 0.104 | ns | 6.899 | 0.010 | ** |
| *w*Aur | 182 | 33.330 | 0.000 | *** | 9.912 | 0.002 | ** | 193.945 | 0.000 | *** | 20.458 | 0.000 | *** | 7.324 | 0.007 | ** | 19.987 | 0.000 | *** |
| *w*Ha | 354 | 35.105 | 0.000 | *** | 4.186 | 0.041 | * | 564.486 | 0.000 | *** | 46.058 | 0.000 | *** | 0.044 | 0.833 | ns | 114.356 | 0.000 | *** |
| *w*Sech | 210 | 27.909 | 0.000 | *** | 19.148 | 0.000 | *** | 397.176 | 0.000 | *** | 14.774 | 0.000 | *** | 0.509 | 0.477 | ns | 11.259 | 0.001 | *** |
| *w*Ha & *w*No | 238 | 25.256 | 0.000 | *** | 6.970 | 0.009 | ** | 195.601 | 0.000 | *** | 164.309 | 0.000 | *** | 21.453 | 0.000 | *** | 62.924 | 0.000 | *** |
| *w*Mau | 146 | 2.735 | 0.100 | ns | 6.848 | 0.010 | ** | 535.908 | 0.000 | *** | 1.675 | 0.197 | ns | 0.220 | 0.639 | ns | 1.740 | 0.189 | ns |

**Supplemental Table S5.** Results from LMMs evaluating the effects of sex and *Wolbachia* on SMR while accounting for body mass.

| | | Sex | | | *Wolbachia* | | | Sex-by-*Wolbachia* | | | Body mass (mg) | | |
|---|---|---|---|---|---|---|---|---|---|---|---|---|---|
| ***Wolbachia* Genotype** | **N** | **F** | **p value** | | **F** | **p value** | | **F** | **p value** | | **F** | **p value** | |
| *w*MelCS | 460 | 0.608 | 0.436 | ns | 0.131 | 0.718 | ns | 1.920 | 0.167 | ns | 282.088 | 0.000 | *** |
| *w*Mel (Maine) | 342 | 35.960 | 0.000 | *** | 7.277 | 0.007 | ** | 0.002 | 0.967 | ns | 139.832 | 0.000 | *** |
| *w*Mel (Panama) | 161 | 1.530 | 0.218 | ns | 3.037 | 0.083 | ns | 0.404 | 0.526 | ns | 27.083 | 0.000 | *** |
| *w*Yak | 164 | 5.229 | 0.024 | * | 0.213 | 0.645 | ns | 0.441 | 0.508 | ns | 92.136 | 0.000 | *** |
| *w*San | 208 | 0.270 | 0.604 | ns | 0.186 | 0.666 | ns | 9.462 | 0.002 | ** | 15.816 | 0.000 | *** |
| *w*Tei | 346 | 0.160 | 0.690 | ns | 0.098 | 0.755 | ns | 0.426 | 0.515 | ns | 22.277 | 0.000 | *** |
| *w*Ri | 129 | 0.526 | 0.470 | ns | 2.576 | 0.111 | ns | 1.487 | 0.225 | ns | 51.688 | 0.000 | *** |
| *w*Suz | 162 | 0.377 | 0.540 | ns | 3.341 | 0.070 | ns | 0.803 | 0.372 | ns | 69.547 | 0.000 | *** |
| *w*Aur | 182 | 10.986 | 0.001 | ** | 1.163 | 0.282 | ns | 0.095 | 0.759 | ns | 157.954 | 0.000 | *** |
| *w*Ha | 354 | 32.319 | 0.000 | *** | 4.788 | 0.029 | * | 0.212 | 0.645 | ns | 117.552 | 0.000 | *** |
| *w*Sech | 210 | 0.524 | 0.470 | ns | 0.001 | 0.970 | ns | 0.009 | 0.923 | ns | 59.01 | 0.000 | *** |
| *w*Ha & *w*No | 238 | 0.113 | 0.737 | ns | 5.796 | 0.017 | * | 0.000 | 0.989 | ns | 59.644 | 0.000 | *** |
| *w*Mau | 146 | 1.769 | 0.186 | ns | 0.026 | 0.873 | ns | 0.462 | 0.498 | ns | 46.056 | 0.000 | *** |

| | | Temperature (C°) | | | Light (lux) | | | Start time | | | Flow rate (ml $min^{-1}$) | | | Water vapor pressure (ppt) | | | Activity (ADS) | | |
|---|---|---|---|---|---|---|---|---|---|---|---|---|---|---|---|---|---|---|---|
| ***Wolbachia* Genotype** | **N** | **F** | **p value** | | **F** | **p value** | | **F** | **p value** | | **F** | **p value** | | **F** | **p value** | | **F** | **p value** | |
| *w*MelCS | 460 | 1.614 | 0.204 | ns | 8.433 | 0.004 | ** | 765.526 | 0.000 | *** | 7.918 | 0.005 | ** | 14.512 | 0.000 | *** | 144.739 | 0.000 | *** |
| *w*Mel (Maine) | 342 | 45.500 | 0.000 | *** | 0.957 | 0.328 | ns | 865.827 | 0.000 | *** | 26.178 | 0.000 | *** | 5.640 | 0.018 | * | 54.583 | 0.000 | *** |
| *w*Mel (Panama) | 161 | 17.740 | 0.000 | *** | 0.062 | 0.804 | ns | 218.883 | 0.000 | *** | 24.623 | 0.000 | *** | 37.504 | 0.000 | *** | 80.620 | 0.000 | *** |
| *w*Yak | 164 | 13.827 | 0.000 | *** | 3.726 | 0.055 | ns | 9.105 | 0.003 | ** | 3.197 | 0.076 | ns | 0.518 | 0.473 | ns | 71.040 | 0.000 | *** |
| *w*San | 208 | 24.112 | 0.000 | *** | 0.824 | 0.365 | ns | 26.204 | 0.000 | *** | 0.494 | 0.483 | ns | 4.653 | 0.032 | * | 80.466 | 0.000 | *** |
| *w*Tei | 346 | 4.610 | 0.032 | * | 8.962 | 0.003 | ** | 23.361 | 0.000 | *** | 0.175 | 0.676 | ns | 4.448 | 0.036 | * | 59.740 | 0.000 | *** |
| *w*Ri | 129 | 2.638 | 0.106 | ns | 0.021 | 0.884 | ns | 40.081 | 0.000 | *** | 6.084 | 0.015 | * | 2.300 | 0.131 | ns | 70.655 | 0.000 | *** |
| *w*Suz | 162 | 22.338 | 0.000 | *** | 4.671 | 0.032 | * | 113.233 | 0.000 | *** | 0.022 | 0.882 | ns | 2.518 | 0.114 | ns | 8.294 | 0.005 | ** |
| *w*Aur | 182 | 29.676 | 0.000 | *** | 10.726 | 0.001 | ** | 197.932 | 0.000 | *** | 1.945 | 0.165 | ns | 0.180 | 0.672 | ns | 35.279 | 0.000 | *** |
| *w*Ha | 354 | 21.338 | 0.000 | *** | 4.137 | 0.042 | * | 568.010 | 0.000 | *** | 113.803 | 0.000 | *** | 1.035 | 0.310 | ns | 136.044 | 0.000 | *** |
| *w*Sech | 210 | 30.049 | 0.000 | *** | 37.064 | 0.000 | *** | 412.464 | 0.000 | *** | 9.689 | 0.002 | ** | 0.114 | 0.736 | ns | 9.842 | 0.002 | ** |
| *w*Ha & *w*No | 238 | 26.746 | 0.000 | *** | 6.594 | 0.011 | * | 201.533 | 0.000 | *** | 131.023 | 0.000 | *** | 24.034 | 0.000 | *** | 70.503 | 0.000 | *** |
| *w*Mau | 146 | 3.600 | 0.059 | ns | 9.232 | 0.003 | ** | 556.934 | 0.000 | *** | 0.006 | 0.936 | ns | 1.101 | 0.295 | ns | 2.212 | 0.138 | ns |

**Supplemental Table S6.** Estimates of mass scaling exponents (slope) with standard error (SE) and confidence intervals for *Wolbachia*-negative and -positive flies for each sex of each *Wolbachia-Drosophila* genotype. Body mass size range represents the order of magnitude difference between the largest and smallest individuals measured (log(*max*) - log(*min*)) for each treatment group.

| Genotype | Sex | *Wolbachia* | N | Size range | Slope | SE | Lower CI | Upper CI |
|---|---|---|---|---|---|---|---|---|
| *w*MelCS | F | - | 118 | 0.456 | 0.669 | 0.067 | 0.538 | 0.8 |
| *w*MelCS | F | + | 116 | 0.361 | 0.75 | 0.08 | 0.593 | 0.907 |
| *w*MelCS | M | - | 110 | 0.385 | 0.598 | 0.081 | 0.439 | 0.757 |
| *w*MelCS | M | + | 116 | 0.35 | 0.789 | 0.108 | 0.578 | 1.001 |
| *w*Mel (Maine) | F | - | 85 | 0.329 | 0.488 | 0.078 | 0.335 | 0.641 |
| *w*Mel (Maine) | F | + | 85 | 0.454 | 0.566 | 0.082 | 0.405 | 0.727 |
| *w*Mel (Maine) | M | - | 86 | 0.334 | 0.519 | 0.084 | 0.354 | 0.685 |
| *w*Mel (Maine) | M | + | 86 | 0.323 | 0.626 | 0.099 | 0.432 | 0.821 |
| *w*Mel (Panama) | F | - | 42 | 0.248 | 0.413 | 0.144 | 0.129 | 0.697 |
| *w*Mel (Panama) | F | + | 41 | 0.234 | 0.425 | 0.169 | 0.092 | 0.759 |
| *w*Mel (Panama) | M | - | 40 | 0.258 | 0.639 | 0.199 | 0.246 | 1.033 |
| *w*Mel (Panama) | M | + | 38 | 0.3 | 0.309 | 0.146 | 0.019 | 0.598 |
| *w*Yak | F | - | 43 | 0.465 | 0.38 | 0.097 | 0.189 | 0.57 |
| *w*Yak | F | + | 40 | 0.477 | 0.523 | 0.092 | 0.342 | 0.704 |
| *w*Yak | M | - | 42 | 0.376 | 0.524 | 0.12 | 0.286 | 0.762 |
| *w*Yak | M | + | 39 | 0.308 | 0.637 | 0.118 | 0.405 | 0.87 |
| *w*San | F | - | 51 | 0.219 | 0.534 | 0.144 | 0.25 | 0.819 |
| *w*San | F | + | 53 | 0.21 | 0.487 | 0.142 | 0.207 | 0.766 |
| *w*San | M | - | 51 | 0.252 | 0.078 | 0.142 | -0.202 | 0.358 |
| *w*San | M | + | 53 | 0.192 | -0.037 | 0.205 | -0.441 | 0.366 |
| *w*Tei | F | - | 97 | 0.63 | 0.334 | 0.106 | 0.124 | 0.543 |
| *w*Tei | F | + | 91 | 0.35 | 0.426 | 0.145 | 0.14 | 0.711 |
| *w*Tei | M | - | 80 | 0.332 | 0.128 | 0.18 | -0.225 | 0.482 |
| *w*Tei | M | + | 78 | 0.345 | 0.47 | 0.176 | 0.123 | 0.817 |
| *w*Ri | F | - | 32 | 0.259 | 0.564 | 0.159 | 0.25 | 0.879 |
| *w*Ri | F | + | 33 | 0.251 | 0.59 | 0.179 | 0.236 | 0.944 |
| *w*Ri | M | - | 31 | 0.347 | 0.711 | 0.129 | 0.456 | 0.966 |
| *w*Ri | M | + | 33 | 0.362 | 0.509 | 0.128 | 0.257 | 0.762 |
| *w*Suz | F | - | 42 | 0.327 | 0.533 | 0.137 | 0.263 | 0.803 |
| *w*Suz | F | + | 46 | 0.461 | 0.696 | 0.101 | 0.496 | 0.896 |
| *w*Suz | M | - | 31 | 0.221 | 0.243 | 0.243 | -0.237 | 0.722 |
| *w*Suz | M | + | 43 | 0.267 | 0.544 | 0.151 | 0.246 | 0.842 |
| *w*Aur | F | - | 45 | 0.445 | 0.89 | 0.075 | 0.743 | 1.037 |
| *w*Aur | F | + | 43 | 0.25 | 0.819 | 0.137 | 0.548 | 1.09 |
| *w*Aur | M | - | 50 | 0.354 | 0.466 | 0.109 | 0.251 | 0.681 |
| *w*Aur | M | + | 44 | 0.296 | 0.407 | 0.13 | 0.151 | 0.664 |
| *w*Ha | F | - | 87 | 0.442 | 0.361 | 0.057 | 0.248 | 0.473 |
| *w*Ha | F | + | 88 | 0.365 | 0.443 | 0.061 | 0.322 | 0.564 |

| | | | | | | | | |
|---|---|---|---|---|---|---|---|---|
| *w*Ha | M | - | 87 | 0.399 | 0.327 | 0.077 | 0.176 | 0.479 |
| *w*Ha | M | + | 92 | 0.365 | 0.312 | 0.075 | 0.164 | 0.46 |
| *w*Sech | F | - | 52 | 0.32 | 0.447 | 0.102 | 0.246 | 0.649 |
| *w*Sech | F | + | 51 | 0.275 | 0.608 | 0.122 | 0.368 | 0.848 |
| *w*Sech | M | - | 52 | 0.228 | 0.378 | 0.121 | 0.138 | 0.617 |
| *w*Sech | M | + | 55 | 0.215 | 0.471 | 0.157 | 0.162 | 0.781 |
| *w*Ha & *w*No | F | - | 57 | 0.314 | 0.735 | 0.125 | 0.488 | 0.982 |
| *w*Ha & *w*No | F | + | 60 | 0.307 | 0.793 | 0.12 | 0.556 | 1.03 |
| *w*Ha & *w*No | M | - | 62 | 0.273 | -0.355 | 0.187 | -0.724 | 0.013 |
| *w*Ha & *w*No | M | + | 59 | 0.29 | 0.682 | 0.154 | 0.378 | 0.986 |
| *w*Mau | F | - | 37 | 0.301 | 0.81 | 0.175 | 0.465 | 1.155 |
| *w*Mau | F | + | 39 | 0.181 | 0.269 | 0.263 | -0.251 | 0.788 |
| *w*Mau | M | - | 34 | 0.334 | 0.764 | 0.169 | 0.43 | 1.098 |
| *w*Mau | M | + | 36 | 0.182 | 0.631 | 0.226 | 0.184 | 1.079 |
